# Functional assignment of Golgi-associated vesicle tethers to specific membrane recycling pathways

**DOI:** 10.64898/2026.05.20.726668

**Authors:** Adam H. Krahn, Natalie Johnson, Jotham Austin, Benjamin S. Glick

## Abstract

During maturation of a Golgi cisterna, multiple vesicular transport pathways recycle resident Golgi proteins. Recycling vesicles are captured by Golgi-associated tethers. To assign individual tethers to specific recycling pathways in *Saccharomyces cerevisiae*, we examined tether arrival and departure using kinetic mapping, and we examined tether function using an ectopic tether localization assay. Those approaches yielded mutually consistent results. Our analysis focused on two coiled coil golgin tethers and the multi-subunit tether GARP. At an intermediate stage of cisternal maturation, the golgin Sgm1 tethers proteins that follow an intra-Golgi recycling pathway dependent on COPI. At a late stage of cisternal maturation, GARP and the golgin Imh1 tether *trans*- Golgi network (TGN) proteins that follow an intra-Golgi recycling pathway dependent on the AP-1 and Ent5 clathrin adaptors. This involvement of GARP in intra-Golgi recycling had not previously been documented. Imh1 also tethers proteins that recycle from prevacuolar endosome compartments to the TGN. Our findings contribute to an integrated model of Golgi membrane traffic.

## Introduction

The Golgi apparatus is a central player in the cellular endomembrane system (Nakano and Saraste, 2026a; Nakano and Saraste, 2026b). This organelle consists of multiple disk-shaped cisternae, which in many eukaryotes are organized into stacks. About 30% of newly synthesized proteins pass through the Golgi, where they encounter a series of processing enzymes that catalyze reactions such as oligosaccharide modification and proteolytic cleavage. In addition, the Golgi is the site of sphingolipid biosynthesis. Newly synthesized proteins and lipids in the Golgi are sorted at the *trans*-Golgi network (TGN) for delivery to other organelles or to the cell exterior.

A conserved feature of the Golgi is that each resident protein is concentrated in a particular subset of the cisternae (Dunphy and Rothman, 1985; Rabouille et al., 1995; Tie et al., 2016; Tojima et al., 2024). The resulting polarized distribution of Golgi enzymes ensures that newly synthesized proteins and lipids are processed efficiently in the appropriate sequence. According to the cisternal maturation model, Golgi polarity is established by membrane traffic pathways that recycle resident Golgi proteins (Glick and Nakano, 2009; Pantazopoulou and Glick, 2019). The core postulate of cisternal maturation is that Golgi cisternae form *de novo* from ER-derived membranes, then progressively mature while carrying the secretory cargoes forward, and then transform into secretory carriers at the TGN stage. Such events can be directly observed in the budding yeast *Saccharomyces cerevisiae*, which contains non-stacked Golgi cisternae that are individually resolvable by fluorescence microscopy (Wooding and Pelham, 1998; Losev et al., 2006; Matsuura-Tokita et al., 2006; Casler et al., 2019). Cisternal maturation is driven by the recycling of resident Golgi proteins from older to younger cisternae. These recycling pathways operate in a defined order, thereby delivering each resident Golgi protein to a cisterna at the appropriate stage of maturation. The distribution of a given resident Golgi protein is determined by the kinetics of its recycling pathway—a protein that recycles early in maturation is present in the early Golgi, whereas a protein that recycles late in maturation is present in the late Golgi (Pantazopoulou and Glick, 2019).

To understand how this dynamic system works, we need to characterize the membrane traffic pathways that recycle resident Golgi proteins. Relevant questions include: (1) How many recycling pathways contribute to cisternal maturation? (2) When during cisternal maturation does each recycling pathway operate? (3) Which traffic machinery components mediate each recycling pathway? (4) Which resident Golgi proteins follow each recycling pathway? *S. cerevisiae* is uniquely powerful for addressing these questions. A key approach is “kinetic mapping” of recycling pathways at the yeast Golgi using 4D fluorescence microscopy, which reveals when various Golgi proteins arrive and depart relative to one another as a cisterna matures (Losev et al., 2006; Papanikou et al., 2015; Tojima et al., 2024). Proteins with similar kinetic signatures often follow the same recycling pathway. However, kinetic mapping is not enough to ascertain whether a given pair of resident Golgi proteins recycle together. We therefore devised a functional assay: recycling vesicles carrying a tagged resident Golgi protein are captured at the yeast bud neck, and a second tagged resident Golgi protein is tested for co-capture with the first protein (Krahn et al., 2026). This combination of kinetic and functional studies enabled us to distinguish between three intra-Golgi recycling pathways that operate at early, intermediate, and late stages of cisternal maturation, plus a fourth recycling pathway that carries certain resident Golgi proteins to prevacuolar endosome (PVE) compartments and back (Casler et al., 2022; Krahn et al., 2026) (Table 1 and Fig. S1).

**Table 1.**
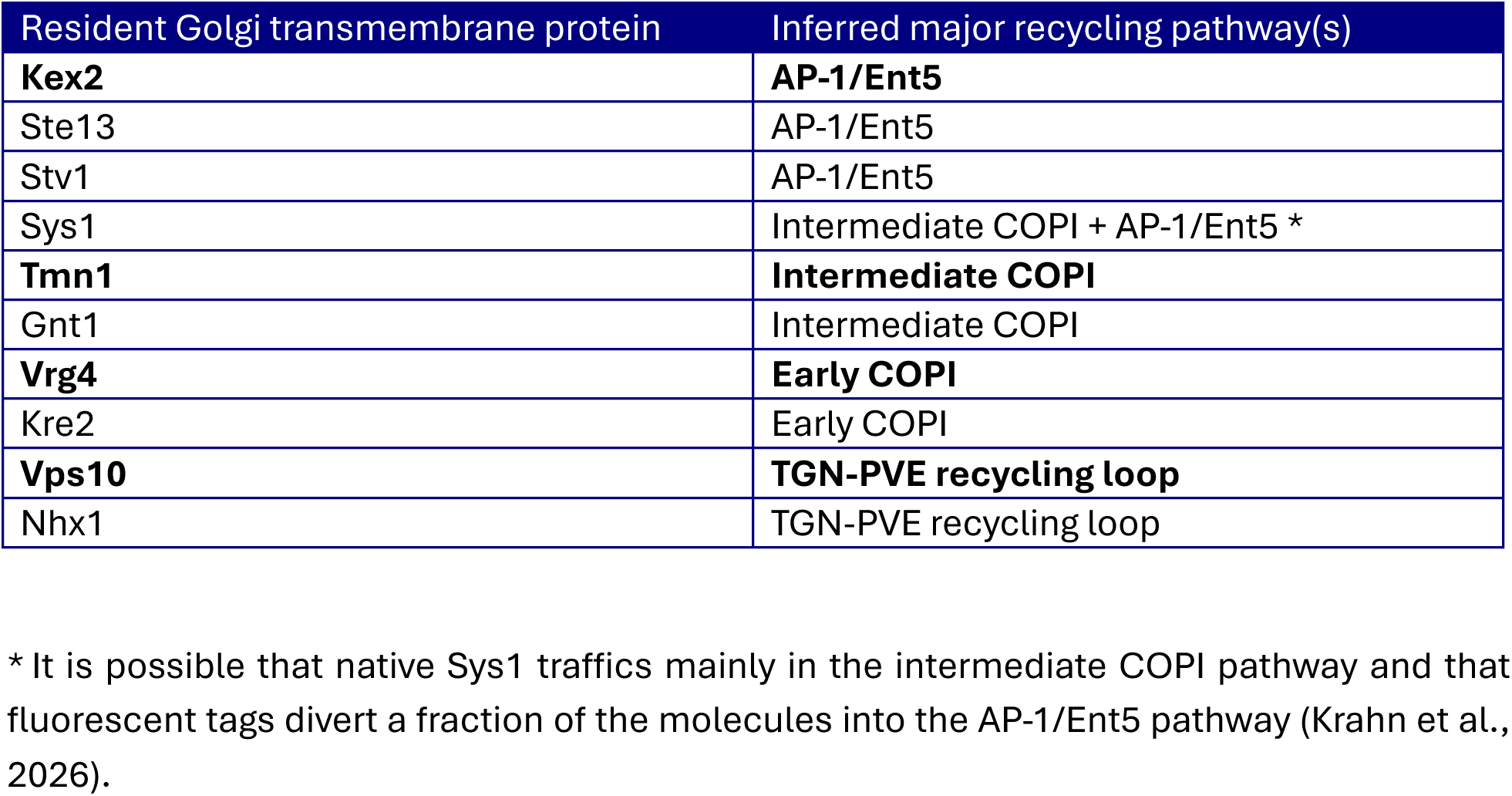
Resident Golgi transmembrane proteins examined in this study. The inferred major recycling pathways are based on our prior work (Casler et al., 2022; Krahn et al., 2026). We have chosen the proteins marked in bold as reference markers for the four pathways examined here. The early COPI pathway was previously designated COPI(b) and the intermediate COPI pathway was previously designated COPI(b’) (Krahn et al., 2026). * It is possible that native Sys1 traffics mainly in the intermediate COPI pathway and that fluorescent tags divert a fraction of the molecules into the AP-1/Ent5 pathway (Krahn et al., 2026).

| Resident Golgi transmembrane protein | Inferred major recycling pathway(s) |
| --- | --- |
| **Kex2** | **AP-1/Ent5** |
| Ste13 | AP-1/Ent5 |
| Stv1 | AP-1/Ent5 |
| Sys1 | Intermediate COPI + AP-1/Ent5 * |
| **Tmn1** | **Intermediate COPI** |
| Gnt1 | Intermediate COPI |
| **Vrg4** | **Early COPI** |
| Kre2 | Early COPI |
| **Vps10** | **TGN-PVE recycling loop** |
| Nhx1 | TGN-PVE recycling loop |

The next step is to identify the molecular components that mediate each of the yeast Golgi recycling pathways. Vesicles generated by the COPI coat are thought to retrieve proteins from the Golgi to the ER and also to recycle proteins within the Golgi (Rabouille and Klumperman, 2005; Barlowe and Miller, 2013), and indeed, we found that the early and intermediate intra-Golgi recycling pathways depend on COPI (Papanikou et al., 2015; Krahn et al., 2026). In our recent publication, those pathways were designated COPI(b) and COPI(b’), respectively (Krahn et al., 2026). The late intra-Golgi recycling pathway depends on the clathrin adaptors AP-1 and Ent5 (Casler et al., 2022; Krahn et al., 2026). For a given recycling pathway, the vesicles need to fuse with cisternae that are at the appropriate stage of maturation. The initial recognition of incoming vesicles by Golgi cisternae is mediated by tethers, which come in two types: coiled coil dimeric tethers termed golgins, and multi-subunit tethers that also help to promote the subsequent steps of vesicle fusion (Gillingham and Munro, 2019; Ungermann and Kümmel, 2019). Most Golgi-associated tethers are peripheral membrane proteins that are recruited with the aid of small GTPases (Panic et al., 2003b; Setty et al., 2003; Fridmann-Sirkis et al., 2004; Ishida and Bonifacino, 2019; Thomas and Fromme, 2020). Thus, the membrane traffic pathways that drive cisternal maturation are coordinated with GTPase cycles (Pantazopoulou and Glick, 2019; Thomas and Fromme, 2020). Studies of ectopically localized tethers in animal cells suggested that different tethers function in distinct recycling pathways (Wong and Munro, 2014; Gillingham and Munro, 2019), but the identities of those recycling pathways were unknown. Here, we expanded on that work by ectopically localizing individual tethers to the yeast bud neck and by determining whether specific resident Golgi proteins were present in the tethered vesicles. Table 1 lists the resident Golgi transmembrane proteins that were used to mark the various recycling pathways.

A challenge for vesicle co-capture and vesicle tethering experiments is that membrane traffic has imperfect fidelity, so a given resident Golgi protein may follow a primary recycling pathway as well as one or more secondary recycling pathways (Glick, 2026; Krahn et al., 2026). With the vesicle co-capture assay that we developed previously, both the Golgi protein used for capture and the Golgi protein being assayed for co-capture can leak into vesicles from secondary recycling pathways, so interpretation of the data requires careful quantification combined with kinetic mapping. The new vesicle tethering assay provides complementary information. We assume that a given ectopically localized tether will recognize vesicles of a particular recycling pathway, in which case the strongest tethering signals should be seen for Golgi proteins that primarily follow that recycling pathway. The results from ectopic tether localization reinforce our earlier conclusions about Golgi-associated recycling pathways and allow us to assign several tethers to specific recycling pathways. These findings enrich our understanding of how vesicular transport maintains the polarized distribution of resident Golgi proteins.

## Results

### Imh1 localizes to Golgi cisternae during a late stage of maturation and can be relocalized to the septin ring

We set out to analyze the golgin Imh1 (Munro and Nichols, 1999; Tsukuda et al., 1999). The first step was kinetic mapping, because the time window when Imh1 is present on maturing cisternae constrains the possible recycling pathways in which Imh1 could operate. To enable kinetic mapping, two or more resident Golgi proteins are fused to fluorescent tags, and then individual cisternae are tracked by time-lapse microscopy to obtain fluorescence traces, and then multiple fluorescence traces are combined to obtain average kinetic signatures (Losev et al., 2006; Johnson and Glick, 2019; Casler et al., 2022). For Imh1, fluorescent tagging is nontrivial because functional domains are present at both termini of the protein. The C-terminal GRIP domain confers Golgi localization (Kjer-Nielsen et al., 1999; Munro and Nichols, 1999; Tsukuda et al., 1999), so we avoided C-terminal tags. By analogy to mammalian GRIP domain golgins, the extreme N-terminal portion of Imh1 is predicted to mediate vesicle binding (Wong et al., 2017), so an N-terminal tag may disrupt tethering but probably will not alter Golgi localization. Based on these considerations, we fused GFP to the N-terminus of Imh1 and expressed this tagged protein as a second copy using the endogenous promoter. The expectation was that the untagged wild-type copy of Imh1 would provide tethering activity while the tagged copy would serve as a reporter for the arrival and departure of this tether.

Our kinetic mapping of Imh1 began with a comparison to the transmembrane protein Sys1, which was previously shown to arrive at cisternae in the COPI-dependent intermediate intra-Golgi recycling pathway (Krahn et al., 2026). The reasoning was as follows. Previous studies showed that Sys1 initiates the biochemical cascade promoting activation and membrane association of the Arl1 GTPase (Behnia et al., 2004; Setty et al., 2004), and that Arl1 then recruits Imh1 to Golgi membranes (Panic et al., 2003b; Setty et al., 2003). Therefore, the prediction was that Imh1 would begin to arrive soon after Sys1. That result was indeed observed (Fig. 1 A). Under our experimental conditions, Imh1 persisted on a maturing cisterna for about a minute on average. Additional kinetic mapping revealed that Imh1 was present when TGN proteins were arriving (Fig. 1 A and Fig. S2 A and Video 1). Specifically, Imh1 was present throughout the arrival phases of Vps10, which recycles to the Golgi from PVE compartments (Marcusson et al., 1994; Cooper and Stevens, 1996), and Kex2, which follows the AP-1/Ent5-dependent intra-Golgi recycling pathway (Casler et al., 2022; Glick, 2026; Krahn et al., 2026). Vps10 and Kex2 began to arrive at approximately the same time, although Vps10 accumulated more rapidly (Fig. 1 A). The presence of Imh1 at the Golgi during the arrival of vesicles carrying Vps10 and of vesicles carrying Kex2 means that Imh1 is a candidate tether for two pathways that recycle TGN proteins.

**Figure 1.**
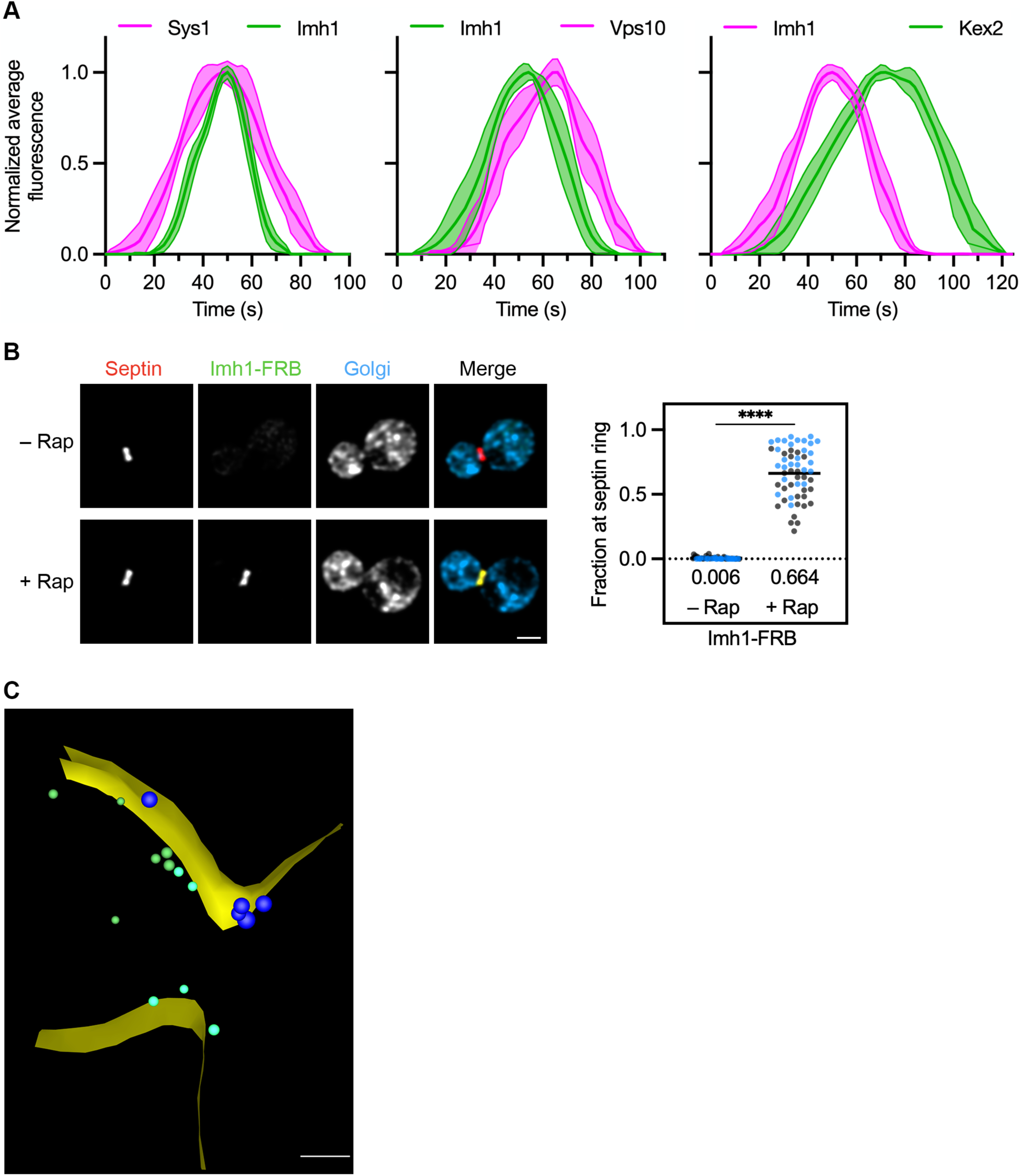
Imh1 is present on maturing cisternae during the arrival of TGN proteins, and a mutant Imh1 can be ectopically localized to the bud neck. **(A)** Golgi maturation kinetics of Imh1 relative to Sys1, Vps10, and Kex2. Green represents a GFP label and magenta represents a HaloTag label. Shown are normalized and averaged traces for 8 individual cisternae for the Sys1/Imh1 comparison, 8 individual cisternae for the Imh1/Vps10 comparison, and 9 individual cisternae for the Imh1/Kex2 comparison. The solid lines are mean values and the shaded areas are 95% confidence intervals. **(B)** Ectopic localization by an FKBP-tagged septin (red) of truncated Imh1 tagged with FRB-GFP (green). HaloTag- labeled Ric1 and Sec7 (blue) marked Golgi cisternae. Representative images are shown. Cells were incubated with vehicle (“– Rap”) or rapamycin (“+ Rap”) for 10 min prior to imaging. Imh1-FRB-GFP levels at the bud neck were quantified by measuring the fraction of the total GFP signal present within a septin mask that had been modified by subtraction of a Golgi mask. The black and blue dots represent two biological replicates. The total numbers of cells examined were 53 (“– Rap”) and 56 (“+ Rap”). Mean values for the combined individual data points from the two biological replicates are displayed as thick horizontal bars and are listed numerically below the plots. ****, significant at P value <0.0001. Multiple regression analysis yielded a P value of <0.0001 that the two replicates were different. **(C)** Cryo-ET of vesicles tethered at the bud neck using FRB-tagged Imh1. A log-phase culture of cells expressing Imh1-FRB and Shs1-FKBP was treated with rapamycin for 10 min followed by cryopreservation and processing for cryo-ET. Shown is the model of a SIRT-reconstructed tomogram from a large budded cell. The full data set is provided in Video 2. A vesicle was counted as putatively tethered if its membrane was no more than 218 nm from a point on the plasma membrane within 200 nm from the center of the bud neck. For the 4 non-rapamycin- treated cells examined, the number of vesicles meeting this criterion ranged from 1 to 7 (mean = 3.0). For the 4 rapamycin-treated cells examined, the number of putatively tethered vesicles ranged from 3 to 11 (mean = 6.0). Cell cortex, yellow; secretory vesicles, blue; putatively tethered non-secretory vesicles, bright green; other non-secretory vesicles, dull green. Scale bar, 250 nm.

To characterize the vesicles tethered by Imh1, we used the rapamycin-inducible heterodimerization system to localize Imh1 ectopically to the yeast bud neck (Haruki et al., 2008; Krahn et al., 2026). In brief, the septin Shs1 was endogenously tagged with mScarlet plus two tandem copies of the FK506-binding protein (FKBP). Then Imh1 was endogenously tagged by replacing the C-terminal GRIP domain, which binds Arl1 to confer Golgi localization (Panic et al., 2003a; Panic et al., 2003b), with GFP plus two copies of the FKBP-rapamycin-binding domain (FRB). The resulting Imh1 fusion protein was largely cytosolic. Despite this displacement of Imh1, the mutant strain had no noticeable growth defect, and TGN proteins such as Kex2 and Ste13 (Table 1) were still visible at the Golgi (see below), although kinetic mapping experiments revealed that Kex2 fluorescence at the Golgi was weaker than normal (data not shown). We proceeded on the assumption that displacement of Imh1 into the cytosol did not severely perturb Golgi traffic and that this mutant strain could therefore be used to study vesicle tethering. Addition of rapamycin relocalized FRB-tagged Imh1 to the septin ring (Fig. 1 B). This relocalization was quantified by measuring the fraction of the total cellular GFP signal that overlapped with the mScarlet-labeled septin ring. The results indicated that in most cells, the majority of the FRB-tagged Imh1 molecules were at the septin ring after a 10-min rapamycin treatment (Fig. 1 B).

We expected that ectopically localized Imh1 would tether and concentrate vesicles at the bud neck, for the following reasons. Transport vesicles presumably travel through the cytosol between membrane compartments. Such vesicles are normally too dispersed to be seen by fluorescence microscopy, but if vesicles carrying a fluorescently tagged Golgi protein collide with septin-bound Imh1 and become tethered, then the resulting concentration of fluorescence should yield a visible signal. This principle was originally established in a study of ectopically localized tethers in mammalian cells (Wong and Munro, 2014). A typical yeast Golgi transmembrane protein recycles about once per minute, so during the 10-min incubation with rapamycin, the entire population of a tagged Golgi protein will experience multiple rounds of vesicular transport, and some fraction of those vesicles should become tethered at the bud neck.

To confirm that ectopic localization of Imh1 led to accumulation of vesicles in the bud neck region, we performed cryo-electron tomography (cryo-ET) of large budded yeast cells (Krahn et al., 2026) by examining lamellae of thickness ∼250-300 nm, representing an estimated 20-40% of the total bud neck volume. Cells expressing FRB-tagged Imh1 were frozen with or without rapamycin treatment under the conditions of the vesicle tethering assay. Although practical considerations limited the output of our cryo-ET pipeline to just a few tomograms per sample, this method was adequate to document vesicle accumulation (Krahn et al., 2026). We were able to examine 4 untreated large budded cells and 4 rapamycin-treated large budded cells. Non-secretory vesicles of diameters ∼35-60 nm located near the septin ring were scored as being putatively tethered if they resided in zones defined by the size of the septin ring, the predicted length of the FKBP-tagged Shs1 septin, and the predicted length of FRB-tagged Imh1 (Krahn et al., 2026) (see Methods). As summarized in Table S1, for the non-rapamycin-treated cells, the number of vesicles meeting this criterion ranged from 1 to 7 (mean = 3.0), whereas for the rapamycin-treated cells, the number of putatively tethered vesicles ranged from 3 to 11 (mean = 6.0). Fig. 1 C and Video 2 show an example of a rapamycin-treated cell for which the lamella included 11 putatively tethered vesicles (marked in bright green). We conclude that ectopically localized Imh1 has vesicle tethering activity.

### Ectopically localized Imh1 tethers vesicles containing TGN proteins

We were now poised to determine whether specific resident Golgi proteins were in the vesicles tethered by Imh1. For this purpose, Imh1 was tagged with FRB alone, and GFP was appended to a transmembrane Golgi protein of interest. The idea was that if the GFP-tagged Golgi protein recycled in vesicles tethered by Imh1, then rapamycin should cause that Golgi protein to accumulate at the septin ring (Fig. 2 A, left). Conversely, if the GFP-tagged Golgi protein recycled in vesicles not tethered by Imh1, then rapamycin should have no visible effect (Fig. 2 A, right). To exclude any signal that might be contributed by Golgi cisternae at the bud neck, we generated a mask by fusing HaloTag to two peripheral membrane proteins: Ric1, which resides at the early Golgi, and Sec7, which resides at the late Golgi (Krahn et al., 2026). HaloTag was conjugated to a far-red dye, and the resulting Golgi mask was subtracted from the GFP fluorescence signal (Krahn et al., 2026). Any remaining GFP signal that overlapped with the septin ring at the bud neck was assumed to represent tethered vesicles. We used a low threshold to create an aggressive Golgi mask, thereby minimizing the background signal from cisternae at the expense of excluding some of the genuine vesicle signal.

**Figure 2.**
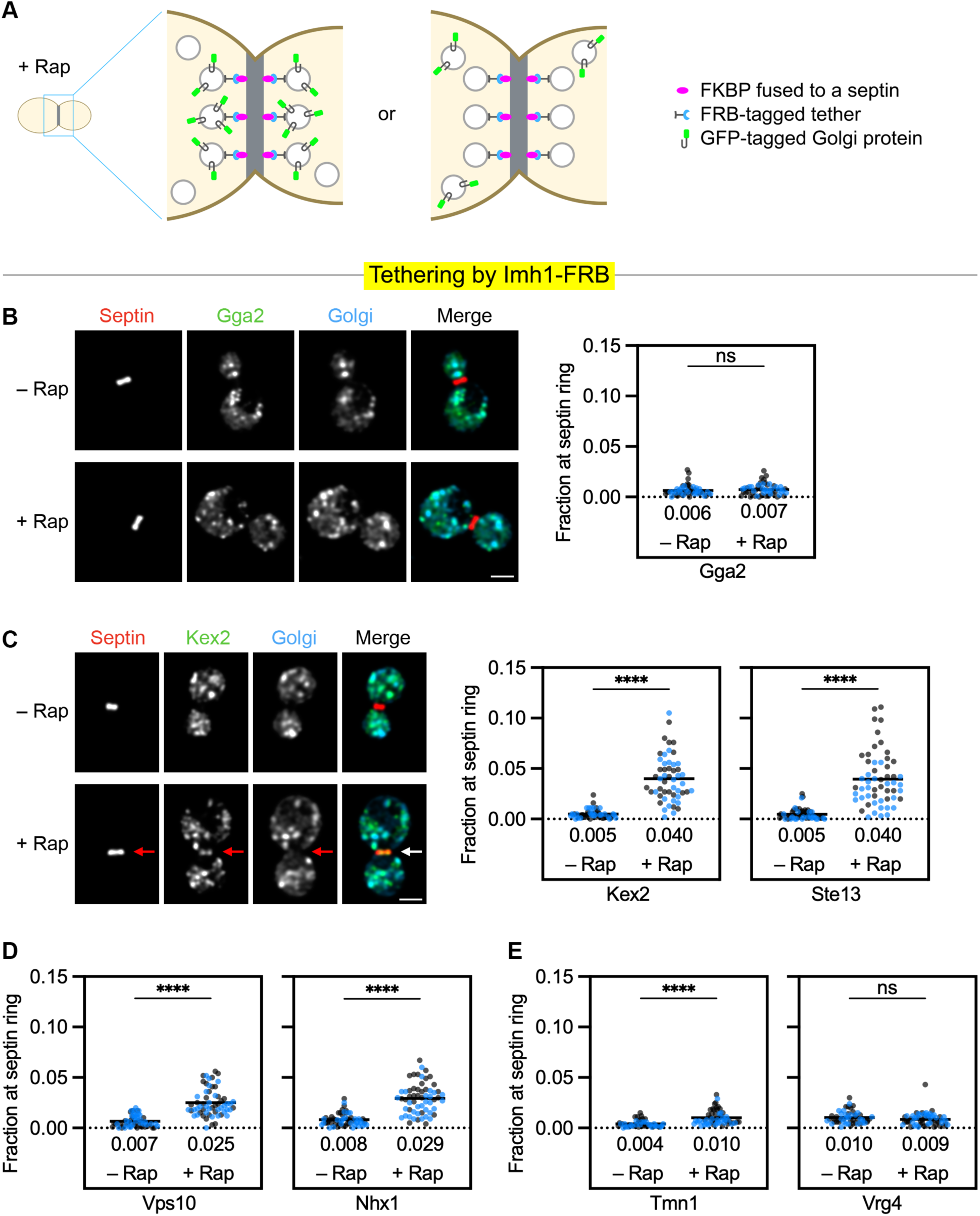
Ectopically localized Imh1 tethers two classes of TGN proteins. For the statistical analyses: ****, significant at P value <0.0001; ns, not significant. **(A)** Diagram of the vesicle tethering assay. A septin at the bud neck is tagged with FKBP (magenta), and a Golgi-associated tether is tagged with FRB (blue) to localize the tether to the bud neck in the presence of rapamycin (“+ Rap”). Meanwhile, a resident Golgi protein is tagged with GFP (green). The ectopically localized tether will recognize transport vesicles. If the GFP-tagged Golgi protein travels in vesicles recognized by the tether, then green fluorescence will be visible at the bud neck (left). Otherwise, no green fluorescence will be visible at the bud neck (right). **(B)** Control experiment showing that ectopic localization of Imh1 to the FKBP-tagged septin (red) did not result in detectable fluorescence at the bud neck from TGN cisternae marked by GFP-tagged Gga2 (green). HaloTag-labeled Ric1 and Sec7 (blue) marked Golgi cisternae. Representative images are shown. Gga2 fluorescence at the bud neck was quantified with or without a 10-min treatment with rapamycin as in Fig. 1 B. The total numbers of cells examined were 52 (“– Rap”) and 51 (“+ Rap”). **(C)** Strong tethering of AP-1/Ent5 cargoes by ectopically localized Imh1. Representative images show that localization of Imh1 to the FKBP-tagged septin (red) resulted in rapamycin-dependent accumulation of GFP-tagged Kex2 (green) at the bud neck. HaloTag-labeled Ric1 and Sec7 (blue) marked Golgi cisternae. Arrows indicate non-Golgi signal at the bud neck. Kex2 fluorescence at the bud neck was quantified with or without a 10-min rapamycin treatment as in Fig. 1 B. A similar quantification was performed for GFP-tagged Ste13. The total numbers of cells examined were 52 (Kex2 “– Rap”), 52 (Kex2 “+ Rap”), 51 (Ste13 “– Rap”), and 55 (Ste13 “+ Rap”). Multiple regression analysis yielded a P value of 0.0006 that the two Ste13 replicates were different. **(D)** Strong tethering of PVE-derived cargoes by ectopically localized Imh1. Fluorescence at the bud neck for GFP-tagged Vps10 or Nhx1 was quantified with or without a 10-min rapamycin treatment as in Fig. 1 B. The total numbers of cells examined were 57 (Vps10 “– Rap”), 60 (Vps10 “+ Rap”), 58 (Nhx1 “– Rap”), and 59 (Nhx1 “+ Rap”). **(E)** Weak or undetectable tethering of COPI cargoes by ectopically localized Imh1. The experiment was performed as in (D) but with GFP-tagged Tmn1 or Vrg4. The total numbers of cells examined were 48 (Tmn1 “– Rap”), 56 (Tmn1 “+ Rap”), 52 (Vrg4 “– Rap”), and 57 (Vrg4 “+ Rap”).

Empirically, tethering by ectopically localized Imh1—and by the other ectopically localized tethers described below—led to trapping at the bud neck of only a small fraction of the GFP signal from a tagged Golgi protein. A possible reason is that the number of tethering sites at the bud neck is limited by the number of septin molecules and/or the number of tether molecules and/or the number of vesicles that can be physically accommodated at the septin ring. Another consideration is that not all of the vesicles will collide with ectopically localized Imh1. Moreover, vesicle tethering is reversible, so the binding affinity may be low, especially when a tether is outside its normal environment. Despite these limitations, we could robustly detect vesicle tethering signals by comparisons with the control incubations lacking rapamycin, and we could distinguish between relatively strong versus weak signals. Because only a small subset of a given vesicle population was trapped at the septin ring during the rapamycin treatment, ectopic tethering likely did not deplete proteins from the Golgi to a degree that would perturb function.

As a control to confirm that subtraction of the Golgi mask eliminated contaminating GFP signal arising from cisternae, we appended a GFP tag to Gga2, a clathrin adaptor found at the TGN (Daboussi et al., 2012; Casler and Glick, 2020). Gga2 is not thought to associate with vesicles that are delivering material to the Golgi (Black and Pelham, 2001), so ectopically localized Imh1 should not alter the distribution of Gga2. Indeed, no vesicle-associated Gga2 was seen at the septin ring after rapamycin addition (Fig. 2 B).

The next step was to test whether ectopically localized Imh1 could tether Kex2 and Ste13, which are TGN proteins that recycle in the AP-1/Ent5 pathway (Casler et al., 2022; Glick, 2026; Krahn et al., 2026). After rapamycin addition, both Kex2 and Ste13 reproducibly accumulated at the septin ring at a level that averaged about 4% of the total fluorescence (Fig. 2 C). Representative images are shown for Kex2. For these and other representative images, we chose examples with tethering signals close to the mean values. As an illustration of an exceptionally strong signal, Fig. S2 B shows a cell with the one of the highest values for tethering of Kex2 by Imh1. These results indicate that Imh1 tethers vesicles from the AP-1/Ent5 pathway.

Ectopically localized Imh1 also tethered Vps10 and Nhx1 (Fig. 2 D and Fig. S2 C), which are TGN proteins that recycle from PVE compartments to the Golgi (Cooper and Stevens, 1996; Kojima et al., 2012; Casler et al., 2022). As described below, follow-up experiments confirmed that Vps10 and Nhx1 follow a different recycling pathway than Kex2 and Ste13. These results indicate that Imh1 tethers vesicles from two pathways that deliver TGN proteins.

Additional experiments examined resident Golgi proteins that recycle in other pathways. Vrg4 follows the COPI-dependent early intra-Golgi recycling pathway (Papanikou et al., 2015; Krahn et al., 2026), and it was not tethered by ectopically localized Imh1 (Fig. 2 E and Fig. S2 D). Weak tethering was seen for Tmn1 (Fig. 2 E), which follows the COPI-dependent intermediate intra-Golgi recycling pathway (Krahn et al., 2026). Based on this finding and previous data (Krahn et al., 2026), we speculate that a fraction of the Tmn1 molecules fail to enter their primary recycling pathway and then are retrieved by the downstream AP-1/Ent5 pathway. A special case is presented by Sys1 (Fig. S2 E). Even though Sys1 is functionally upstream of Imh1 and begins to arrive before Imh1, ectopically localized Imh1 showed moderately strong tethering of Sys1. At face value, this result might seem surprising. However, our previous work revealed that a substantial fraction of GFP-tagged Sys1 molecules partition into the AP-1/Ent5 pathway (Krahn et al., 2026), so the AP-1/Ent5 vesicles tethered by Imh1 are expected to contain Sys1. Thus, the various results obtained with kinetic mapping and ectopic tether localization suggest a coherent interpretation: Imh1 tethers vesicles that arrive downstream of the COPI-dependent recycling pathways.

### Two classes of TGN proteins follow distinct recycling pathways

After seeing that Imh1 tethered all four of the TGN proteins examined, we sought to verify that Kex2 and Ste13 follow a different recycling pathway than Vps10 and Nhx1. This point is important because Kex2 and Ste13 were long thought to recycle together with Vps10 between PVE compartments and the Golgi (Voos and Stevens, 1998; Bowers and Stevens, 2005; Bean et al., 2017). Instead, our data suggest that the primary recycling route of Kex2 and Ste13 is the intra-Golgi AP-1/Ent5 pathway, and that this pathway was originally overlooked due to occasional missorting of Kex2 and Ste13 to PVE compartments (Glick, 2026). A key piece of evidence for two distinct pathways is that deleting AP-1 plus Ent5 perturbs the traffic of Kex2 and Ste13 but not of Vps10 or Nhx1 (Casler et al., 2022). Moreover, Kex2 and Ste13 localize almost exclusively to the Golgi whereas Vps10 and Nhx1 localize to both PVE compartments and the Golgi (Kojima et al., 2012; Chi et al., 2014; Day et al., 2018; Casler et al., 2022). Here, we sought additional ways to characterize the different pathways.

One approach was to compare kinetic signatures. All four TGN proteins began to arrive at about the same time, slightly before the TGN marker Sec7 (Day et al., 2018; Casler et al., 2022) (Fig. 3 A). However, the departure times varied. Vps10 departed much earlier than Kex2, and Nhx1 departed somewhat earlier than Kex2 (Fig. 3 A). By contrast, Ste13 departed synchronously with Kex2 (Fig. 3 B). Synchronous departure may be a general property of the AP-1/Ent5 pathway because Stv1, another cargo of that pathway (Casler et al., 2022; Krahn et al., 2026), also departed together with Kex2 (Fig. 3 B). Thus, Vps10 and Nhx1 depart from maturing cisternae earlier than Kex2 and Ste13 and Stv1, consistent with the existence of separate traffic pathways.

**Figure 3.**
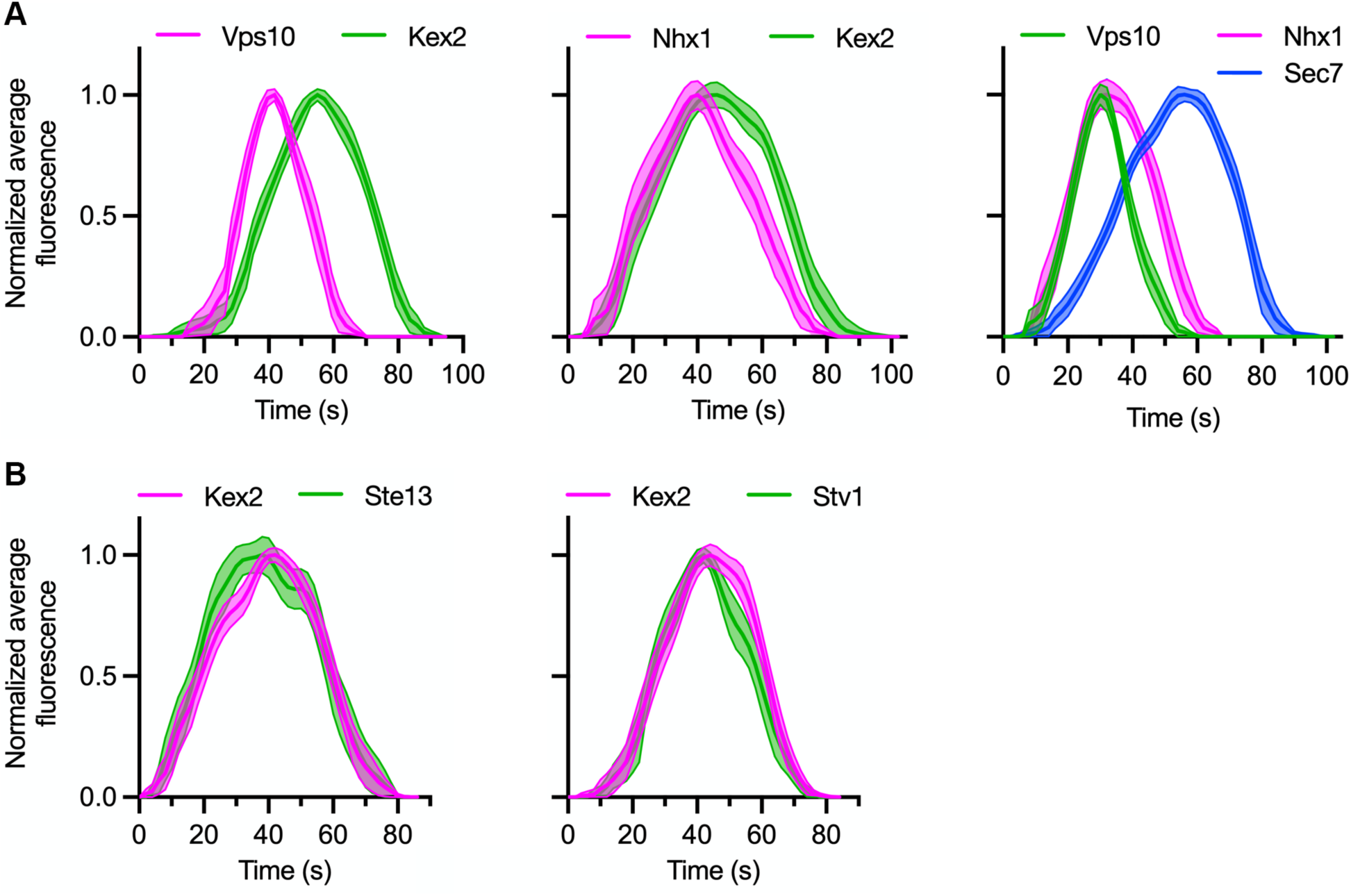
Kinetic mapping distinguishes two classes of TGN proteins. Green represents a GFP label, magenta represents a HaloTag label, and blue represents an mScarlet label. Shown are normalized and averaged traces. **(A)** Golgi maturation kinetics of Vps10 and Nhx1 relative to Kex2 and relative to each other. Because Vps10 and Nhx1 are present both in the TGN and in PVE compartments, the comparison between these two proteins also included Sec7 to identify TGN structures. Shown are normalized and averaged traces for 17 individual cisternae for the Vps10/Kex2 comparison, 17 individual cisternae for the Nhx1/Kex2 comparison, and 16 individual cisternae for the Vps10/Nhx1/Sec7 comparison. **(B)** Golgi maturation kinetics of Kex2 relative to the other AP-1/Ent5 cargoes Ste13 and Stv1. Shown are normalized and averaged traces for 18 individual cisternae for the Kex2/Ste13 comparison and 16 individual cisternae for the Kex2/Stv1 comparison.

A second approach employed our original vesicle co-capture assay (Krahn et al., 2026) to determine which proteins travel together with Vps10. When Vps10 was tagged with both FRB and GFP, an average of 16% of the fusion protein molecules could be captured at the bud neck after 5 min of rapamycin treatment (Fig. 4 A). For this experiment, the Golgi mask was extended by fusing HaloTag to Vps8, which marks PVE compartments (Arlt et al., 2015; Day et al., 2018). The resulting Golgi/PVE mask was subtracted before making the measurements, so the Vps10 fluorescence signal at the bud neck presumably represented captured vesicles that were traveling between PVE compartments and the Golgi. Indeed, when Vps10 was tagged with FRB alone, cryo-ET confirmed that rapamycin treatment resulted in the accumulation of vesicles in the bud neck region (Fig. S3 A and Video 3). In a control co-capture experiment, the GFP-tagged PVE marker Hse1 (Bilodeau et al., 2002; Henne et al., 2011) showed only very weak rapamycin-dependent co-capture with Vps10 (Fig. 4 B and Fig. S4 A), indicating that the Golgi/PVE mask removed most of the fluorescence signal from PVE compartments at the bud neck. As expected, Nhx1 was efficiently co-captured with Vps10 (Fig. 4 C), indicating that the two proteins were in the same vesicles. Ste13 was not detectably co-captured (Fig. 4 C). Kex2 showed some co-capture with Vps10, consistent with low-level missorting of Kex2 to PVE compartments, but this signal was much weaker than the Nhx1 signal (Fig. 4 C). These results support the conclusion that the primary recycling pathway of Kex2 and Ste13 differs from that of Vps10 and Nhx1.

**Figure 4.**
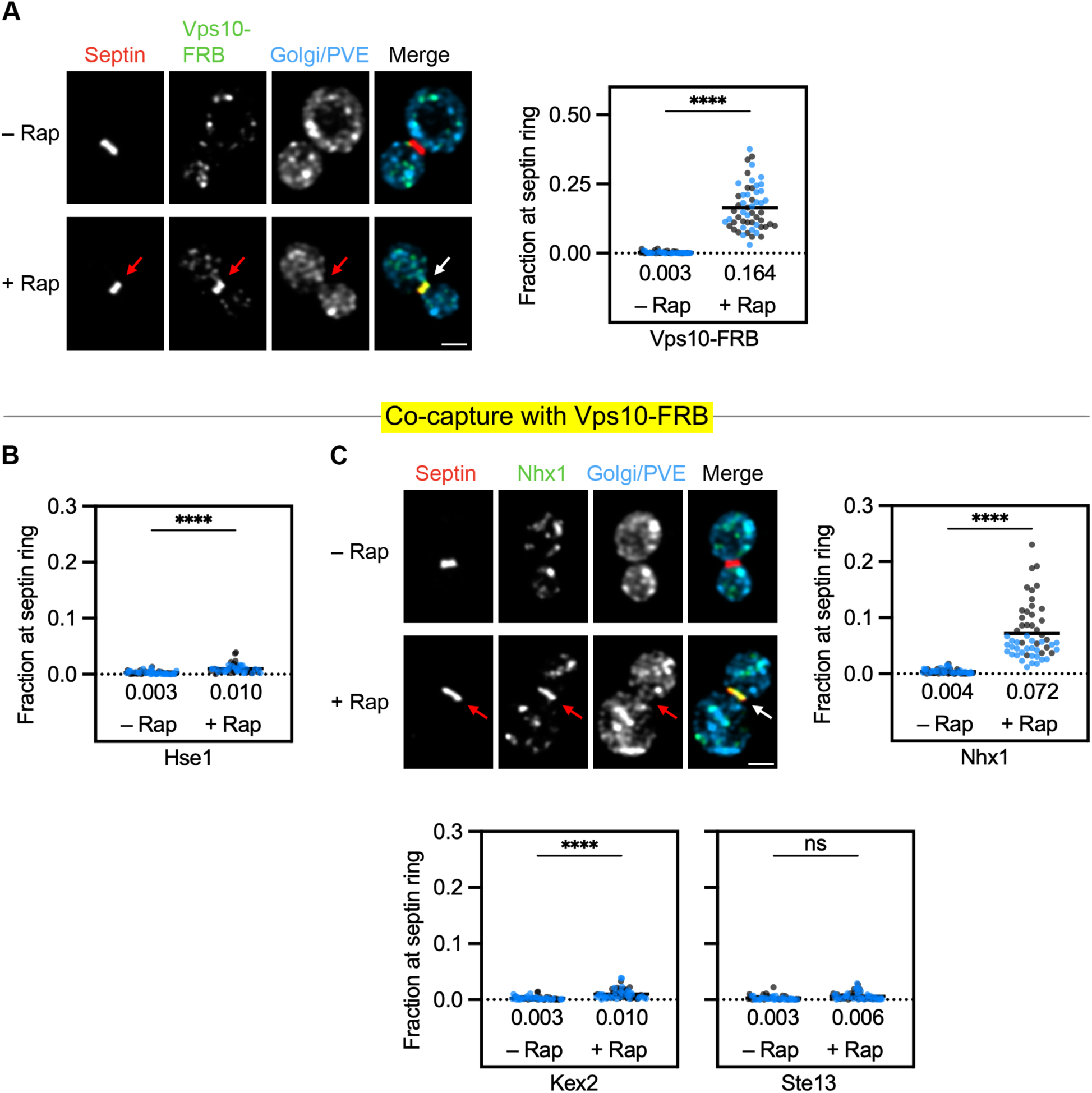
A vesicle co-capture assay distinguishes two classes of TGN proteins. For the statistical analyses: ****, significant at P value <0.0001; ns, not significant. **(A)** Rapamycin- dependent capture by an FKBP-tagged septin (red) of Vps10-FRB-GFP (green). HaloTag- labeled Ric1, Sec7, and Vps8 (blue) marked Golgi cisternae and PVE compartments. Representative images are shown. Arrows indicate non-Golgi and non-PVE signal at the bud neck after treatment for 5 min with rapamycin (“+ Rap”). Vps10-FRB fluorescence at the bud neck was quantified with or without a 5-min treatment with rapamycin as in Fig. 1 B. The total numbers of cells examined were 54 (“– Rap”) and 55 (“+ Rap”). **(B)** Control experiment showing that capture of Vps10-FRB-containing vesicles at the bud neck resulted in minimal co-capture of GFP-tagged Hse1, which marks PVE compartments. Quantification was performed as in (A). The total numbers of cells examined were 57 (“– Rap”) and 55 (“+ Rap”). **(C)** Rapamycin-dependent co-capture with Vps10-FRB of GFP-tagged Nhx1, and weak or undetectable co-capture with Vps10-FRB of GFP-tagged Kex2 or Ste13. Quantification was performed as in (A). The total numbers of cells examined were 56 (Nhx1 “– Rap”), 54 (Nhx1 “+ Rap”), 54 (Kex2 “– Rap”), 60 (Kex2 “+ Rap”), 53 (Ste13 “– Rap”), and 61 (Ste13 “+ Rap”). Multiple regression analysis yielded a P value of <0.0001 that the two Nhx1 replicates were different. Representative images are shown for Nhx1. Arrows indicate non-Golgi and non- PVE signal at the bud neck.

We also performed a complementary experiment by tagging Kex2 with FRB (Fig. S4 B). For reference, Fig. S4 B includes our previously published data showing that Ste13 was co-captured with Kex2 (Krahn et al., 2026). Vps10 and Nhx1 were also co-captured with Kex2, although less strongly. We suspect that the signals from Vps10 and Nhx1 reflect occasional missorting of Kex2 to PVE compartments followed by retrieval. Evidence for such missorting includes weak Kex2 signals in the PVE compartments of wild-type cells (Day et al., 2018) and strong Kex2 signals in the PVE compartments of mutant cells defective in retrieving Kex2 to the Golgi (Nothwehr and Hindes, 1997; Voos and Stevens, 1998; Glick, 2026). Even a single missorted Kex2 molecule in a vesicle traveling between a PVE compartment and the Golgi could lead to rapamycin-dependent capture of that vesicle, together with its cargo of Vps10 and Nhx1. This argument illustrates that individual vesicle co-capture and vesicle tethering assays have caveats, which we try to address by evaluating the data in a holistic way.

### GARP localizes to Golgi cisternae during a late stage of maturation and can be relocalized to the septin ring

The Golgi-associated retrograde protein (GARP) complex is a multi-subunit tether that localizes to the late Golgi (Bonifacino and Hierro, 2011; Khakurel et al., 2021), but the precise time when GARP is present during cisternal maturation has not been reported. GARP is a heterotetramer composed of the proteins Vps51, Vps52, Vps53, and Vps54 (Siniossoglou and Pelham, 2002; Conibear et al., 2003; Reggiori et al., 2003). We kinetically mapped Vps52 relative to Vps10, Kex2, and Imh1 (Fig. 5 A). GARP arrived and departed almost synchronously with Vps10, although GARP initially accumulated somewhat more slowly than Vps10. GARP began to arrive at about the same time as Kex2 and accumulated slightly faster than Kex2. GARP departed earlier than Kex2, but it was present throughout the arrival phase of Kex2. Finally, GARP arrived and departed about 10-20 sec after Imh1, resulting in partial kinetic overlap between these two tethers.

**Figure 5.**
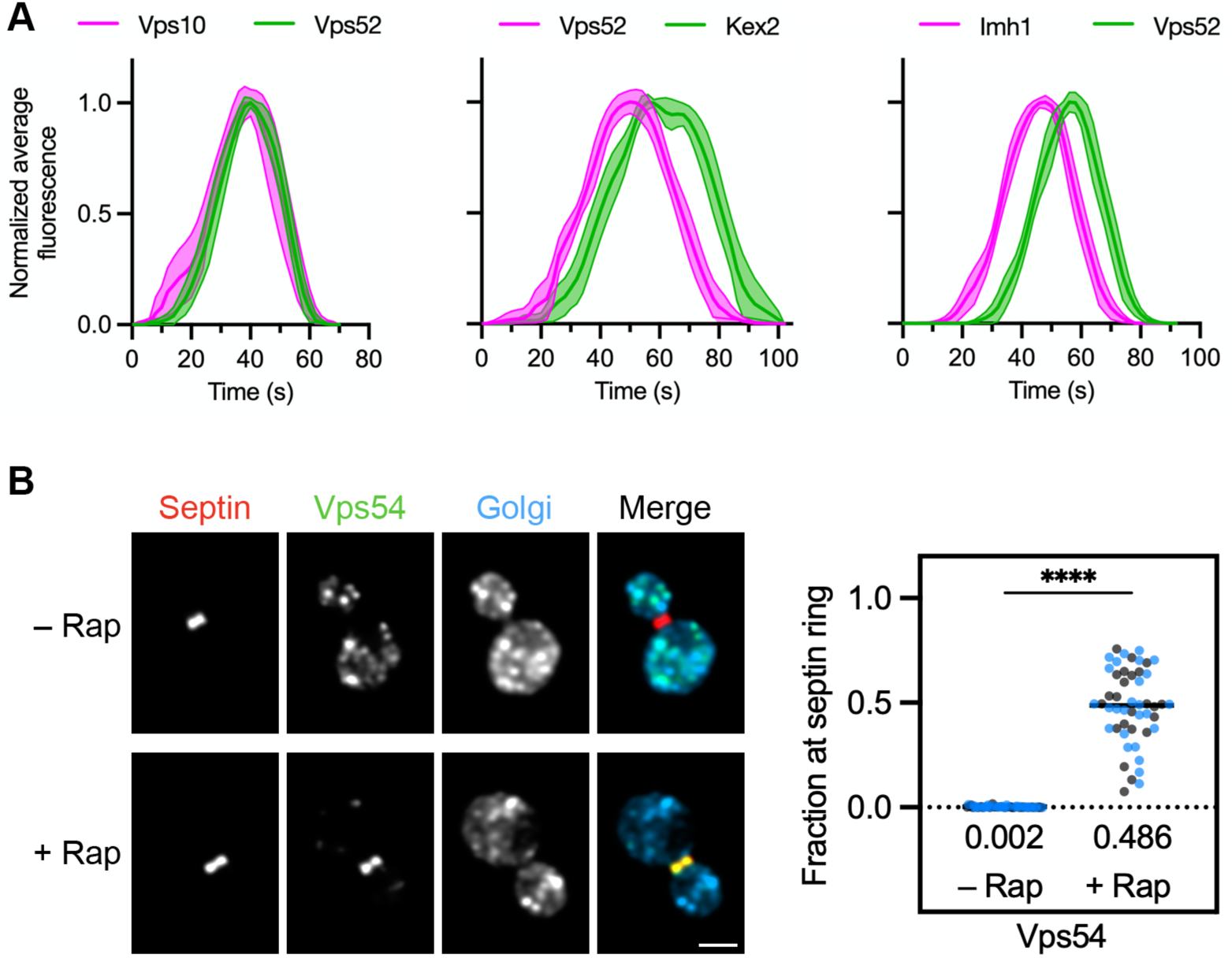
GARP is present on cisternae during a late stage of maturation, and FRB-tagged GARP can be ectopically localized to the bud neck. (**A)** Golgi maturation kinetics of the GARP subunit Vps52 relative to Vps10, Kex2, and Imh1. Green represents a GFP label and magenta represents a HaloTag label. Shown are normalized and averaged traces for 9 individual cisternae for the Vps10/Vps52 comparison, 9 individual cisternae for the Vps52/Kex2 comparison, and 10 individual cisternae for the Imh1/Vps52 comparison. **(B)** Ectopic localization by an FKBP-tagged septin (red) of a GARP complex containing Vps52- FRB and Vps54-GFP (green). HaloTag-labeled Ric1 and Sec7 (blue) marked Golgi cisternae. Representative images are shown. Vps54-GFP levels at the bud neck were quantified as in Fig. 1 B. The total numbers of cells examined were 54 (“– Rap”) and 49 (“+ Rap”). ****, significant at P value <0.0001.

To achieve ectopic localization of GARP, Vps52 was tagged with FRB. Relocalization of GARP was assayed by tagging Vps54 with GFP. Treatment with rapamycin for 10 min eliminated most of the punctate Golgi signal and redistributed GARP to the septin ring (Fig. 5 B). This ectopic localization of GARP led to accumulation of vesicles in the bud neck region as visualized by cryo-ET (Fig. S3 B and Video 4). We conclude that ectopically localized GARP has vesicle tethering activity.

### Ectopically localized GARP tethers vesicles carrying cargoes of the AP-1/Ent5 pathway

The next question was whether ectopically localized GARP could tether one or both of the vesicle types that recycle TGN proteins. A control experiment showed no rapamycin-dependent redistribution of Gga2 by GARP (Fig. S5 A), confirming that the Golgi mask eliminated contaminating fluorescence from TGN cisternae. GARP tethered significant amounts of Kex2 and Ste13, which follow the AP-1/Ent5 pathway (Fig. 6 A). Because GARP is likely to tether endocytic vesicles that fuse directly with the Golgi, and because Kex2 contains an endocytosis signal (Tan et al., 1996), we wondered if tethering of Kex2 by GARP might reflect occasional exocytosis followed by endocytosis. This explanation is unlikely because tethering of Kex2 by GARP was unaffected by CK-666, which potently inhibits endocytosis (Burke et al., 2014; Casler et al., 2022) (Fig. 6 A). We infer that GARP can recognize AP-1/Ent5 vesicles.

**Figure 6.**
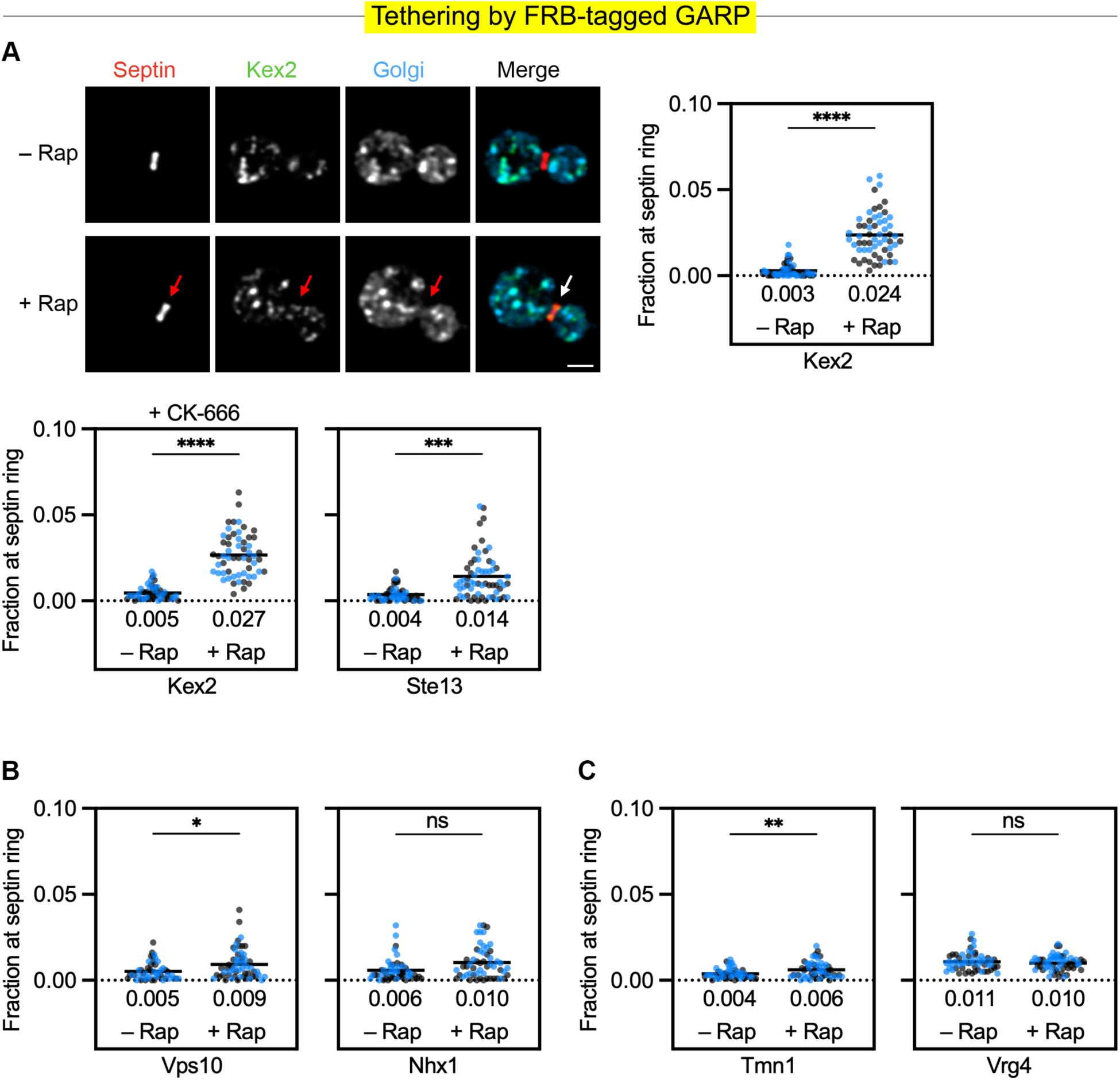
Ectopically localized GARP tethers AP-1/Ent5 cargoes. For the statistical analyses: ****, significant at P value <0.0001; ***, significant at P value = 0.0001 (Ste13); **, significant at P value = 0.0097 (Tmn1); *, significant at P value = 0.0405 (Vps10); ns, not significant. **(A)** Moderate to strong tethering of AP-1/Ent5 cargoes by ectopically localized GARP. The Vps52 subunit of GARP was tagged with FRB. Representative images show that ectopic localization of GARP to the FKBP-tagged septin (red) resulted in rapamycin- dependent accumulation of GFP-tagged Kex2 (green) at the bud neck. HaloTag-labeled Ric1 and Sec7 (blue) marked Golgi cisternae. Arrows indicate non-Golgi signal at the bud neck. Kex2 fluorescence at the bud neck was quantified with or without a 10-min rapamycin treatment as in Fig. 1 B. Similar quantifications were performed for GFP-tagged Kex2 in cells treated for 15 min with CK-666 and for GFP-tagged Ste13. The total numbers of cells examined were 57 (Kex2 “– Rap”), 58 (Kex2 “+ Rap”), 53 (Kex2 + CK-666 “– Rap”), 57 (Kex2 + CK-666 “+ Rap”), 52 (Ste13 “– Rap”), and 61 (Ste13 “+ Rap”). **(B)** Minimal tethering of PVE- derived cargoes by ectopically localized GARP. Fluorescence at the bud neck for GFP-tagged Vps10 or Nhx1 was quantified with or without a 10-min rapamycin treatment as in Fig. 1 B. The total numbers of cells examined were 49 (Vps10 “– Rap”), 57 (Vps10 “+ Rap”), 51 (Nhx1 “– Rap”), and 54 (Nhx1 “+ Rap”). **(C)** Weak or undetectable tethering of COPI cargoes by ectopically localized GARP. The experiment was performed as in (B) but with GFP-tagged Tmn1 or Vrg4. The total numbers of cells examined were 50 (Tmn1 “– Rap”), 53 (Tmn1 “+ Rap”), 57 (Vrg4 “– Rap”), and 58 (Vrg4 “+ Rap”).

The mechanism that recruits GARP to the Golgi is poorly understood, and we wondered whether soluble GARP might bind in the cytosol to AP-1/Ent5 vesicles, which would then deliver GARP to Golgi cisternae. This idea was tested by using a vesicle co-capture assay with Kex2-FRB to see if GARP would bind to Kex2-containing AP-1/Ent5 vesicles at the septin ring. No such binding was seen (Fig. S5 B), suggesting that GARP is recruited to Golgi membranes rather than to vesicles. The most likely interpretation is that GARP normally associates with Golgi membranes and then tethers AP-1/Ent5 vesicles.

With Vps10 and Nhx1, which travel from PVE compartments to the Golgi, tethering by GARP was minimal (Fig. 6 B and Fig. S5 C). This result is at odds with previous interpretations about the function of yeast GARP (Conibear and Stevens, 2000; Conibear et al., 2003; Bonifacino and Hierro, 2011). However, those studies were based on analysis of GARP knockout strains, and we observed that loss of GARP disrupts both late and early Golgi organization, presumably due to a combination of direct and indirect effects (data not shown). Taken together, our results suggest that Imh1 and GARP cooperate to tether AP-1/Ent5 vesicles, but that Imh1 can tether PVE-derived vesicles without assistance from GARP.

Because GARP arrives downstream of cargoes that follow the COPI-dependent intra-Golgi recycling pathways, GARP was not expected to tether cargoes of those pathways. No tethering was seen with Vrg4, which follows the early recycling pathway (Fig. 6 C). A very low level of tethering was seen with Tmn1, which follows the intermediate recycling pathway (Fig. 6 C), but this weak signal resembles the weak tethering of Tmn1 by Imh1 (see Fig. 2 E) and it probably reflects occasional leakage of Tmn1 into the AP-1/Ent5 pathway. We conclude that GARP tethers vesicles from the AP-1/Ent5 pathway but not from the other pathways examined in this study.

### Sgm1 localizes to Golgi cisternae during an early stage of maturation and can be relocalized to the septin ring

The golgin Sgm1 is an effector of the Rab GTPase Ypt6, which is present on Golgi cisternae during early and intermediate stages of maturation (Siniossoglou and Pelham, 2001; Fridmann-Sirkis et al., 2004; Suda et al., 2013) (Fig. 7 A). Based on the same reasoning that was used for kinetic mapping of Imh1 (see above), HaloTag was fused to the N-terminus of Sgm1, and tagged Sgm1 was expressed as a second copy in a strain that also expressed untagged Sgm1. As expected, Sgm1 accumulated on Golgi cisternae at about the same time as Ypt6 (Fig. 7 A). A surprising result is that Sgm1 departed well before Ypt6 (Fig. 7 A). The departure of Sgm1 coincided with that of Vrg4 (Fig. 7 B), perhaps indicating that Golgi association of Sgm1 requires a second factor that recycles synchronously with Vrg4. In any case, Sgm1 was present on cisternae during an early stage of maturation.

**Figure 7.**
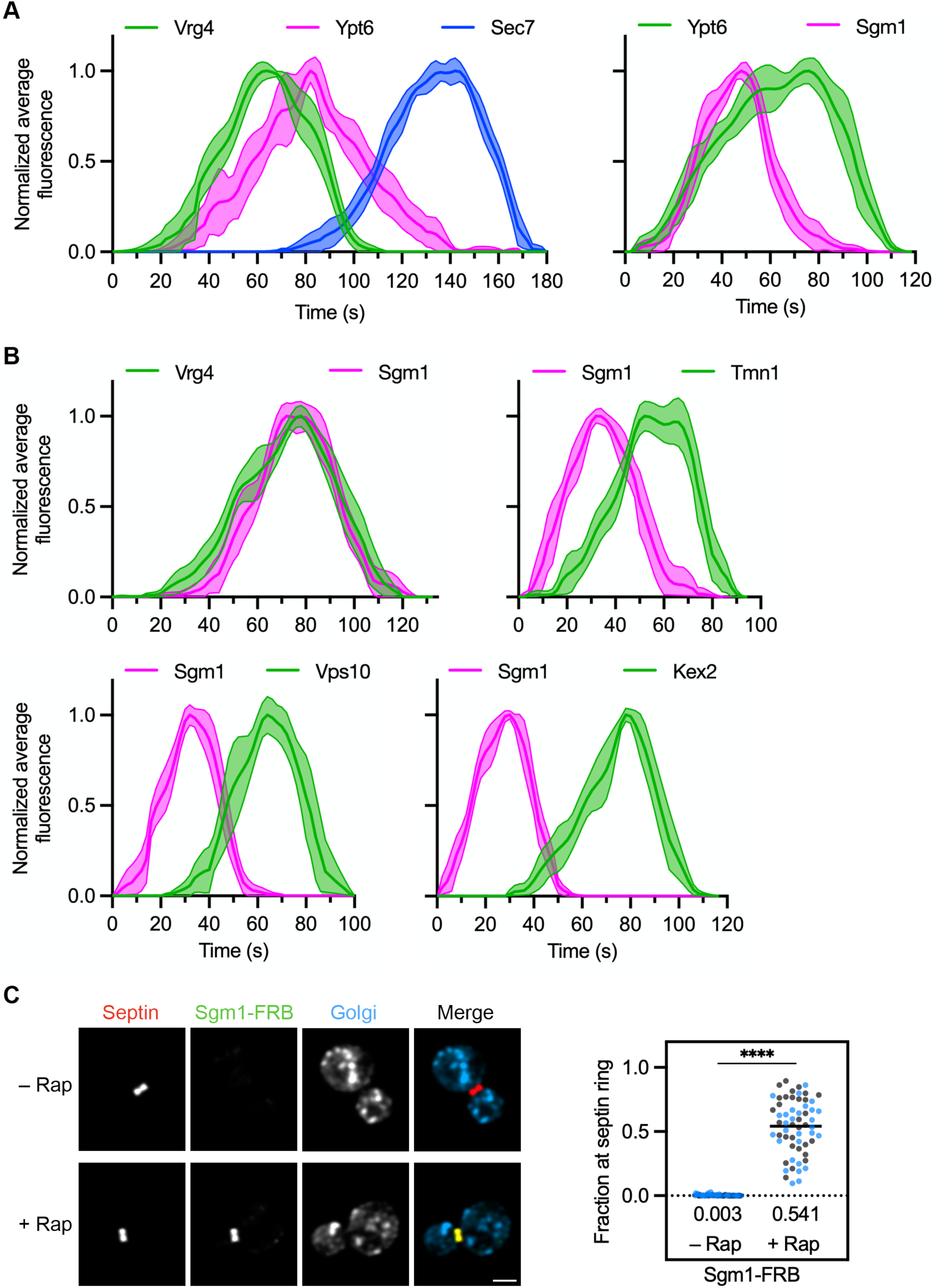
Sgm1 is present on maturing cisternae during the arrival of vesicles of the intermediate intra-Golgi recycling pathway, and a mutant Sgm1 can be ectopically localized to the bud neck. **(A)** Golgi maturation kinetics of Ypt6 relative to the early and late Golgi markers Vrg4 and Sec7, respectively, and relative to Sgm1. Green represents a GFP label, magenta represents a HaloTag label, and blue represents an mScarlet label. Shown are normalized and averaged traces for 8 individual cisternae for the Vrg4/Ypt6/Sec7 comparison and 8 individual cisternae for the Ypt6/Sgm1 comparison. **(B**) Golgi maturation kinetics of Sgm1 relative to Vrg4, Tmn1, Vps10, and Kex2. The display format is as in (A). Shown are normalized and averaged traces for 12 individual cisternae for the Vrg4/Sgm1 comparison, 8 individual cisternae for the Sgm1/Tmn1 comparison, 8 individual cisternae for the Sgm1/Vps10 comparison, and 8 individual cisternae for the Sgm1/Kex2 comparison. **(C)** Ectopic localization by an FKBP-tagged septin (red) of a truncated Sgm1 tagged with FRB-GFP (green). HaloTag-labeled Ric1 and Sec7 (blue) marked Golgi cisternae. Representative images are shown. Sgm1-FRB-GFP levels at the bud neck were quantified as in Fig. 1 B. The total numbers of cells examined were 54 (“– Rap”) and 60 (“+ Rap”). ****, significant at P value <0.0001.

We compared the kinetic signature of Sgm1 with the arrival phases of Vrg4, Tmn1, Vps10, and Kex2 (Fig. 7 B). Accumulation of Sgm1 lagged behind that of Vrg4 (Fig. 7 B), making it unlikely that Sgm1 tethers vesicles of the COPI-dependent early intra-Golgi recycling pathway. Sgm1 was present throughout the arrival phase of Tmn1 (Fig. 7 B), so Sgm1 is a candidate tether for the COPI-dependent intermediate intra-Golgi recycling pathway. Sgm1 was largely absent during the arrival phases of Vps10 and Kex2 (Fig. 7 B), so Sgm1 is not a candidate tether for PVE-derived vesicles or AP-1/Ent5 vesicles.

For the tethering assay, Sgm1 was endogenously tagged to enable ectopic localization by replacing the C-terminal Ypt6-binding domain (Fridmann-Sirkis et al., 2004) with GFP plus two copies of FRB. The resulting Sgm1 fusion protein was largely cytosolic. Despite this displacement of Sgm1, the mutant strain had no noticeable growth defect, and there was no obvious change in the patterns of the Golgi proteins that we examined (see below). We proceeded on the assumption that displacement of Sgm1 into the cytosol did not severely perturb Golgi traffic and that this mutant strain could therefore be used to study vesicle tethering. Addition of rapamycin relocalized FRB-tagged Sgm1 to the septin ring (Fig. 7 C). This ectopic localization of Sgm1 led to accumulation of vesicles in the bud neck region as visualized by cryo-ET (Fig. S3 C and Video 5). We conclude that ectopically localized Sgm1 has vesicle tethering activity.

### Ectopically localized Sgm1 tethers vesicles of the COPI-dependent intermediate intra-Golgi recycling pathway

The next step was to tag Sgm1 with FRB alone and to tag a transmembrane Golgi protein of interest with GFP. After addition of rapamycin, ectopically localized Sgm1 tethered vesicles containing Tmn1, Gnt1, and Sys1 (Fig. 8 A), all of which follow the COPI-dependent intermediate intra-Golgi recycling pathway (Krahn et al., 2026). For unknown reasons, tethering of Sys1 was particularly strong. No tethering was seen for Vps10 or Kex2 (Fig. 8 B). In the case of Vrg4 and Kre2, which follow the COPI-dependent early intra-Golgi recycling pathway, very weak tethering was seen (Fig. 8 B), probably because those proteins occasionally fail to recycle in the COPI-dependent early pathway and then undergo salvage in the COPI-dependent intermediate pathway (Krahn et al., 2026). These results fit with the kinetic mapping, and they suggest that Sgm1 uniquely tethers vesicles of the COPI-dependent intermediate intra-Golgi recycling pathway.

**Figure 8.**
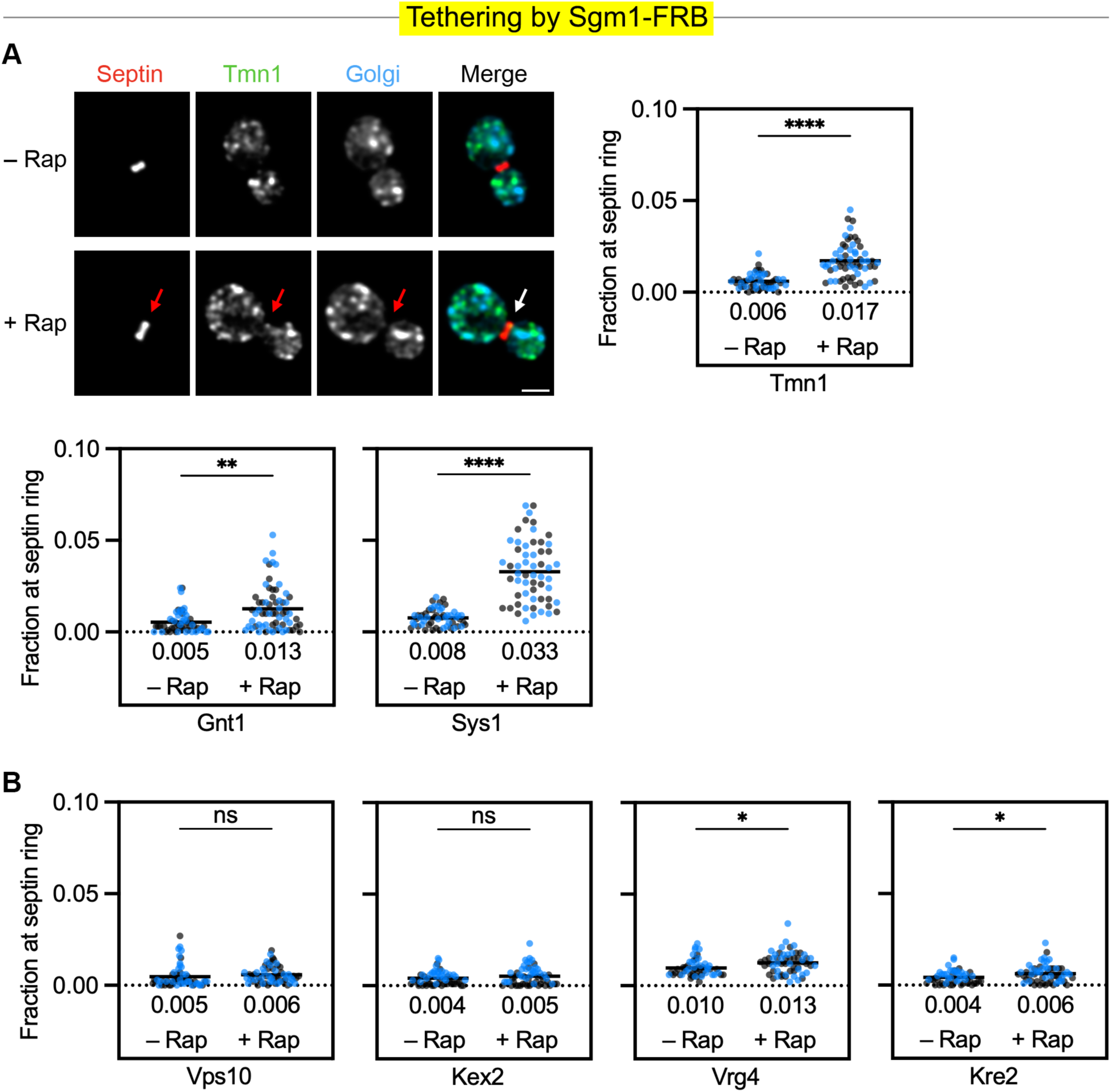
Ectopically localized Sgm1 tethers proteins that follow the intermediate intra-Golgi recycling pathway. For the statistical analyses: ****, significant at P value <0.0001; **, significant at P value = 0.0052 (Gnt1); *, significant at P value = 0.010 (Vrg4) or 0.018 (Kre2); ns, not significant. **(A)** Moderate to strong tethering of three cargoes by ectopically localized Sgm1. Representative images show that ectopic localization of Sgm1 to the FKBP-tagged septin (red) resulted in rapamycin-dependent accumulation of GFP-tagged Tmn1 (green) at the bud neck. HaloTag-labeled Ric1 and Sec7 (blue) marked Golgi cisternae. Arrows indicate non-Golgi signal at the bud neck. Tmn1 fluorescence at the bud neck was quantified with or without a 10-min rapamycin treatment as in Fig. 1 B. Similar quantifications were performed for GFP-tagged Gnt1 and Sys1. The total numbers of cells examined were 53 (Tmn1 “– Rap”), 61 (Tmn1 “+ Rap”), 53 (Gnt1 “– Rap”), 64 (Gnt1 “+ Rap”), 54 (Sys1 “– Rap”), and 57 (Sys1 “+ Rap”). **(B)** Weak or undetectable tethering by ectopically localized Sgm1 of cargoes that follow other recycling pathways. Fluorescence at the bud neck for GFP-tagged Vps10, Kex2, Vrg4, or Kre2 was quantified with or without a 10-min rapamycin treatment as in Fig. 1 B. The total numbers of cells examined were 58 (Vps10 “– Rap”), 59 (Vps10 “+ Rap”), 56 (Kex2 “+ Rap”), 58 (Kex2 “– Rap”), 54 (Vrg4 “– Rap”), 58 (Vrg4 “+ Rap”), 52 (Kre2 “– Rap”), and 58 (Kre2 “+ Rap”).

## Discussion

Our objective was to assign vesicle tethers to membrane recycling pathways at the yeast Golgi. The obvious approach would have been to knock out individual tethers, but such experiments have shortcomings that depend on the strengths of the knockout phenotypes. Loss of a tether might have only a mild effect that is not reliably detectable with our assays. This concern seems to apply to the golgins examined here. Alternatively, loss of a tether might substantially compromise a given pathway, in which case additional pathways could be indirectly affected. This concern seems to apply to the multi-subunit tether GARP. Because of these factors, our preliminary knockout data were largely uninformative, and we chose not to pursue that line of investigation.

A more promising strategy is to have a positive readout for the function of a tether. The starting point was our development of an assay in which Golgi-derived vesicles are captured at the yeast bud neck via binding of a tagged Golgi protein present in the vesicles (Krahn et al., 2026). Here, we modified this assay by ectopically localizing a tether to the bud neck and then assessing whether particular Golgi proteins become trapped at the bud neck. This method was inspired by earlier work on animal Golgi tethers from the Munro lab (Wong and Munro, 2014; Gillingham and Munro, 2016). By quantifying the tethering of Golgi proteins that follow specific recycling pathways, we can determine which classes of vesicles are recognized by each tether.

A limitation of vesicle tethering assays is that membrane traffic systems have imperfect fidelity—Golgi proteins that follow a primary recycling pathway will undergo occasional missorting into secondary pathways, often followed by retrieval. Missorting and retrieval are well documented for resident ER proteins (Barlowe and Miller, 2013), but such processes likely also play a key role in the Golgi. This phenomenon is illustrated by the Kex2 processing protease in the yeast TGN (Fuller et al., 1988). Kex2 was previously thought to cycle between PVE compartments and the Golgi together with the vacuolar hydrolase receptor Vps10 (Voos and Stevens, 1998; Bowers and Stevens, 2005; Bean et al., 2017), yet unlike Vps10, Kex2 shows very low steady-state localization to PVE compartments (Day et al., 2018). We have argued that Kex2 normally follows the AP-1/Ent5-dependent late intra-Golgi recycling pathway, and that Kex2 sometimes undergoes missorting to PVE compartments followed by retrieval to the Golgi (Glick, 2026). The experimental consequence is that when Kex2 is tagged with FRB, rapamycin will cause capture of AP-1/Ent5 vesicles plus less efficient capture of vesicles traveling between the Golgi and PVE compartments. Similarly, when a tether for the AP-1/Ent5 pathway is ectopically localized, the tethered vesicles will contain high levels of proteins such as Kex2 that normally follow the AP-1/Ent5 pathway plus low levels of other proteins that are occasionally missorted into that pathway. Therefore, we must try to distinguish between the relatively strong vesicle tethering signals from primary recycling pathways versus the weaker signals from secondary pathways.

Vesicle tethering data are complemented by kinetic mapping, in two ways. First, the kinetic signature of a resident Golgi protein provides information about the primary recycling pathway(s) for that protein (Papanikou et al., 2015; Day et al., 2018; Casler et al., 2022; Krahn et al., 2026). Second, if a vesicle tether operates in a particular recycling pathway, then the tether must be present during the arrival phase for resident Golgi proteins traveling in that pathway. By combining kinetic data with vesicle tethering data, we have assigned recycling pathways to two golgin tethers as well as the multi-subunit tether GARP.

The first golgin we examined was Imh1, the sole GRIP domain protein in yeast (Kjer- Nielsen et al., 1999; Munro and Nichols, 1999; Tsukuda et al., 1999). Association of Imh1 with the Golgi is triggered by arrival of the transmembrane protein Sys1, which initiates a biochemical cascade that recruits and activates the small GTPase Arl1 (Panic et al., 2003b; Setty et al., 2003; Behnia et al., 2004; Setty et al., 2004). We showed previously that Sys1 recycles in the COPI-dependent intermediate intra-Golgi recycling pathway, although a fraction of the Sys1 molecules partition instead into the AP-1/Ent5-dependent late intra-Golgi recycling pathway (Krahn et al., 2026). As expected, Imh1 associates with a maturing cisterna soon after the first wave of Sys1 arrival. That timing puts Imh1 in place during the arrival of both AP-1/Ent5 vesicles and PVE-derived vesicles. Indeed, ectopically localized Imh1 robustly tethers multiple TGN proteins: Kex2 and Ste13, which recycle in the AP-1/Ent5 pathway, as well as Vps10 and Nhx1, which recycle between PVE compartments and the TGN (Day et al., 2018; Casler et al., 2022; Glick, 2026; Krahn et al., 2026).

This dual tethering by Imh1 prompted us to seek concrete evidence that different classes of TGN proteins follow distinct recycling pathways. We examined the four TGN proteins listed above plus Stv1, which recycles in the AP-1/Ent5 pathway (Casler et al., 2022; Krahn et al., 2026). In wild-type cells, only Vps10 and Nhx1 show clear concentration in PVE compartments in addition to the TGN (Kojima et al., 2012; Chi et al., 2014; Day et al., 2018), suggesting that those two proteins follow a different recycling pathway than the other three. Support for that interpretation came from our earlier analysis of strains carrying deletions of both Ent5 and the AP-1 subunit Apl4 (Casler et al., 2022; Krahn et al., 2026). In *apl4*Δ *ent5*Δ mutant cells, Kex2, Ste13, and Stv1 escape from the Golgi to the plasma membrane whereas Vps10 and Nhx1 traffic normally. Further insight came from kinetic mapping. Although all five TGN proteins begin to arrive at about the same time, the departure times vary. We find that Kex2, Ste13, and Stv1 depart from a maturing cisterna synchronously, whereas Vps10 and Nhx1 depart earlier. This timing fits with observations that the GGA and Ent3 clathrin adaptors that mediate Golgi-to-PVE traffic arrive earlier than the AP-1 and Ent5 adaptors that mediate intra-Golgi recycling (Daboussi et al., 2012; Casler and Glick, 2020; Casler et al., 2022). The combined data support the idea that Vps10 and Nhx1 recycle between PVE compartments and the Golgi in a pathway that involves the GGA adaptors and possibly Ent3, whereas Kex2, Ste13, and Stv1 recycle within the Golgi in a pathway that involves AP-1 and Ent5.

To explore this issue further, we used a vesicle co-capture assay. When Vps10 is tagged with FRB, Nhx1 is co-captured efficiently, whereas Kex2 is only very weakly co-captured and Ste13 is not detectably co-captured. This result is a further indication that Vps10 and Nhx1 follow a different primary recycling pathway than Kex2 and Ste13. Because Imh1 tethers vesicles carrying all four of these TGN proteins, Imh1 evidently recognizes two distinct types of vesicles (Fig. S1).

A golgin such as Imh1 is expected to cooperate with a multi-subunit tether such as GARP (Chen et al., 2019), but our analysis of GARP yields a more nuanced picture. Imh1 and GARP show only a partial kinetic overlap, with Imh1 arriving and departing earlier than GARP, suggesting only a limited functional partnership between the two tethers. The best-established role of GARP is in the tethering and fusion of vesicles from the endocytic system (Bonifacino and Hierro, 2011). Yeast GARP recognizes endocytic vesicles that are targeted to the Golgi (see below). Although yeast GARP has also been proposed to tether PVE-to-Golgi vesicles that carry proteins such as Vps10 (Conibear and Stevens, 2000; Conibear et al., 2003), the evidence for that idea came from GARP knockout strains, which show widespread defects at both early and late Golgi stages (data not shown). Our kinetic mapping suggests that accumulation of GARP at the Golgi lags slightly behind that of Vps10, making it less likely that GARP acts as a tether for Vps10-containing vesicles. Indeed, ectopically localized GARP shows minimal tethering of Vps10 and Nhx1. By contrast, for TGN proteins that recycle in AP-1/Ent5 vesicles, the results are different: GARP arrives simultaneously with or slightly earlier than Kex2, and ectopically localized GARP tethers Kex2 and Ste13, suggesting that GARP functions in AP-1/Ent5-dependent intra-Golgi recycling. We conclude that tethering of AP-1/Ent5 vesicles involves both Imh1 and GARP while tethering of PVE-to-Golgi vesicles involves Imh1 but not GARP (Fig. S1).

The second golgin we examined was Sgm1, the ortholog of mammalian TMF (Siniossoglou and Pelham, 2001; Fridmann-Sirkis et al., 2004). Sgm1 is an effector of the Rab GTPase Ypt6, which is present during an intermediate stage of maturation (Suda et al., 2013). We confirmed that Sgm1 arrives around the same time as Ypt6. Unexpectedly, Sgm1 departs before Ypt6, possibly indicating that a second component is needed to retain Sgm1 at the Golgi membrane. Kinetic data reveal that Sgm1 is present during arrival of vesicles of the COPI-dependent intermediate intra-Golgi recycling pathway but not during arrival of vesicles of the other recycling pathways. These kinetics are consistent with the finding that ectopically localized Sgm1 shows relatively strong tethering of proteins such as Tmn1 and Sys1 that follow the intermediate pathway. For Vrg4 and Kre2, which follow the COPI-dependent early intra-Golgi recycling pathway, a weak signal is seen, presumably due to occasional leakage of those proteins into the intermediate pathway (Krahn et al., 2026). Thus, Sgm1 may uniquely recognize vesicles of the intermediate pathway (Fig. S1).

Our overall conclusion is that vesicle tethers at the yeast Golgi can be assigned to specific membrane traffic pathways (Fig. S1). Imh1 and GARP seem to operate in more than one pathway while Sgm1 seems to operate in a single pathway. The various tethers and membrane traffic pathways are functionally interconnected. For example, the golgin Sgm1 recognizes vesicles of the COPI-dependent intermediate intra-Golgi recycling pathway, which delivers the transmembrane protein Sys1, which triggers a biochemical cascade that recruits the golgin Imh1, which recognizes vesicles of the AP-1/Ent5 and PVE-to-Golgi pathways. Future experiments will flesh out these links by examining additional tethers, including the golgin Rud3, which localizes to the early Golgi (Kim, 2003; Gillingham et al., 2004), and the multi-subunit tether COG, which likely functions in multiple membrane traffic pathways at the Golgi (Blackburn et al., 2019). The ultimate goal is to elucidate the molecular logic circuit that regulates Golgi maturation by switching membrane traffic pathways on and off and by triggering the association and dissociation of GTPases, tethers, and other peripheral membrane proteins (Pantazopoulou and Glick, 2019; Thomas and Fromme, 2020).

The findings presented here extend work from the Munro lab, who ectopically localized golgins in mammalian cells (Wong and Munro, 2014; Gillingham and Munro, 2016). Mammalian golgins probably function similarly to their yeast counterparts, based on the following evidence. First, like Imh1, the mammalian GRIP domain-containing golgin-97 and golgin-245 tether TGN proteins that either recycle within the TGN in an AP-1-dependent pathway or recycle from endosomes (Shin et al., 2017; Navarro Negredo et al., 2018; Cattin-Ortolá et al., 2024). Second, like Sgm1, mammalian TMF tethers proteins that reside in an intermediate part of the Golgi and that probably recycle with the aid of COPI (Wong and Munro, 2014). With the yeast system, the major step forward is that we have characterized recycling pathways for resident Golgi proteins and so we can assign vesicle tethers to well-defined pathways.

The conserved multi-subunit GARP complex can now be added to the list of tethers studied by ectopic localization. GARP was previously implicated in tethering vesicles from the endocytic system (Bonifacino and Hierro, 2011). In yeast, endocytic vesicles fuse directly with maturing Golgi cisternae, which also serve as early endosomes (Day et al., 2018; Glick, 2026), and GARP operates in that endocytic pathway (A. Pantazopoulou and A. Krahn, in preparation). By contrast, GARP does not seem to tether PVE-to-Golgi vesicles. Extrapolating to mammalian cells, GARP might promote the tethering and fusion of early endosome-derived vesicles but not of late endosome-derived vesicles. We also uncovered a novel role for GARP in tethering intra-Golgi AP-1/Ent5 vesicles. Future work will seek to clarify how multi-subunit tethers cooperate with golgins to ensure specificity of vesicle targeting during Golgi membrane traffic.

## Materials and methods

### Yeast growth and transformation

The strains used in this study were derivatives of JK9-3da (*leu2-3,112 ura3-52 rme1 trp1 his4*; Kunz et al., 1993) with the *pdr1Δ pdr3Δ* mutations to facilitate HaloTag labeling (Barrero et al., 2016; Casler et al., 2022). We previously described the construction of rapamycin-resistant strains carrying the *fpr1Δ* and *TOR1-1* mutations (Papanikou et al., 2015). If a Golgi protein was luminally tagged with GFP or HaloTag, the strain carried the *vps10-104* mutation to prevent missorting to the vacuole (Fitzgerald and Glick, 2014; C. Lee-Smith, in preparation). Prior to experimental analysis, yeast cells were grown in nonfluorescent minimal glucose medium (NSD) (Bevis et al., 2002) in baffled flasks at 23°C with shaking at 200 rpm.

Yeast genes were tagged by chromosomal gene replacement using the pop-in/pop-out method (Rothstein, 1991; Rossanese et al., 1999). In most cases, a clean gene replacement was performed. However, cells expressing an N-terminally tagged golgin tether or N-terminally tagged Ypt6 were from a pop-in strain that contained both the untagged, functional wild-type protein and the tagged protein, both of which were expressed from the native promoter. The plasmids used for those manipulations were constructed with the aid of SnapGene software (Dotmatics) and the Saccharomyces Genome Database (Wong et al., 2023), and they have been archived at Addgene together with annotated SnapGene files that describe how the plasmids were linearized for integration.

### Reagents

HaloTag ligands of JFX_646_ and JFX_650_ (Grimm et al., 2021), generously provided by Luke Lavis (Janelia Research Campus, Ashburn, Virginia, USA), were dissolved in anhydrous DMSO (Invitrogen, catalog no. D12345) at a concentration of 1 mM, and single-use aliquots were stored at -80°C. These stock solutions were diluted 1000-fold to label cells at a final dye concentration of 1 µM. Rapamycin (LC Laboratories, catalog no. R-5000) was prepared as a 1 mg/ml solution in 90% ethanol, 10% Tween 20, and single-use aliquots were stored at -20°C. This stock solution was diluted 100-fold to treat cells at a final rapamycin concentration of 10 µg/ml. CK-666 (Sigma, catalog no. SML0006) was dissolved in anhydrous DMSO to a concentration of 50 mM, and single-use aliquots were stored at 4°C. This stock solution was diluted 500-fold to treat cells at a final CK-666 concentration of 100 µM.

### HaloTag labeling

Labeling of HaloTag constructs (Casler et al., 2022) was performed as follows. HaloTag ligand of JFX_646_ or JFX_650_ was diluted by adding 1 µl of a 1 mM DMSO stock solution to 300 µl of NSD. The resulting solution was cleared of precipitate by spinning at 17,000xg (13,000 rpm) in a microcentrifuge for 5 min. Then the supernatant was added to 700 µl of mid-log phase yeast culture to give a final concentration of 1 µM dye, and the cells were incubated for 30-60 min at 23°C with shaking. For live cell imaging experiments, excess dye was removed by filtration and washing with a 0.22-µm syringe filter (Millipore; catalog no. SLGV004SL). The washed cells on the filter were resuspended in NSD and adhered to a coverglass-bottom dish coated with concanavalin A (ConA) (Johnson and Glick, 2019). Confocal videos were then immediately acquired as described below.

### Live cell fluorescence microscopy and analysis

For 4D confocal microscopy, yeast cells were attached to ConA-coated coverglass-bottom dishes that were filled with NSD (Johnson and Glick, 2019). Imaging was performed at 23°C using a Leica Stellaris confocal microscope equipped with a 1.4-NA/63x oil objective. Imaging of cell volumes was performed with 40-80 nm pixels, a 0.30 µm z-step interval, and 20-30 optical sections. z-stacks were obtained at intervals of 2 s. For two-color imaging, both channels used HydX detectors. For three-color imaging, the green channel used a HydS detector and the red and far-red channels used HydX detectors. Image acquisition was performed using the Leica LAS X software.

Confocal movies of Golgi cisternal maturation events were deconvolved using Huygens software (SVI) and quantified with custom ImageJ plugins as previously described (Johnson and Glick, 2019), except that Huygens software was employed for time-based correction of photobleaching prior to deconvolution. Although the kinetic traces showed some cisterna-to-cisterna variation, the basic patterns were highly reproducible between different cisternae in the same or different cells and between movies generated in different imaging sessions. For a typical analysis, we tracked one or two cisternae each from 8-18 cells visualized in one or two imaging sessions. When kinetic traces from a single cisterna were plotted, the fluorescence trace in each channel was normalized to the average of the three highest values. When averaged kinetic traces from 8-18 cisternae were plotted, the normalization and averaging were accomplished using custom ImageJ plugins (Casler et al., 2022). The plots of averaged traces show solid lines for mean values plus shaded areas for 95% confidence intervals.

### Vesicle co-capture and vesicle tethering assays

These methods were adapted from our prior study (Krahn et al., 2026). Cells grown overnight at 23°C to an OD_600_ of ∼0.5 were stained with 1 µM JFX_650_ HaloTag ligand as described above, except that the final volume was 500 µl. After growth with shaking for 1 h at 23°C, either a 100-fold dilution of 90% ethanol, 10% Tween 20 was added as a vehicle control or rapamycin was added to a final concentration of 10 µg/ml. For the vesicle co-capture assays employing Vps10-FRB and Kex2-FRB, the cells were incubated with shaking for an additional 5 min, whereas for the vesicle tethering assays employing FRB-tagged tethers, the cells were incubated with shaking for an additional 10 min. Where indicated, the cells were treated for 15 min with CK-666 prior to the 10-min rapamycin treatment. Then fixation was performed by adding 250 µl of the culture while vortexing to 750 µl of 1.33-fold concentrated fixative to give final concentrations of 1% paraformaldehyde plus 0.1% glutaraldehyde (Electron Microscopy Sciences, diluted from freshly opened vials), 1 mM MgCl_2_, 50 mM potassium phosphate, pH 6.5. After fixation on ice for 1 h, the cells were washed twice by centrifuging for 2 min at 1500xg (4000 rpm) and resuspending in 500 µl of phosphate-buffered saline (PBS). Finally, the cells were centrifuged once again and resuspended in 20 µl PBS. Within 30 h of fixation, the fixed cells were compressed under coverslips and imaged with a Leica Stellaris confocal microscope using a 1.4-NA/63x oil objective with 40 nm pixels, a 0.20 µm z-step interval, and 21 optical sections. Cells were chosen for analysis if they were a mother-daughter cell pair with a joined septin ring, and if Golgi cisternae were largely absent from the bud neck region as determined by viewing the Golgi marker channel. The cell images were deconvolved using Huygens software to smooth the punctate structures and remove background fluorescence (Johnson and Glick, 2019). A set of 20-30 cells was then quantified as described below.

For quantification of vesicle co-capture or vesicle tethering, a deconvolved confocal image stack of a cell was processed with a custom ImageJ plugin termed “Quantify Overlap”, available with source code from https://github.com/bsglicker/4D-Image-Analysis. With this plugin, the user chooses a region of interest (ROI), and the ROI measurements are summed for each image in the z-stack. As used here, the plugin creates a mask from the signal in the red channel and then compares the green signal within the mask to the total green signal in the ROI. The red channel mask can be modified by creating a second mask from the signal in the blue channel and then subtracting the blue channel mask from the red channel mask. Threshold levels are chosen empirically by the user for the red and blue channel masks, from a set of threshold options generated by a custom algorithm. For our purposes, the red threshold was set to the “Lower” level, and the blue threshold was set to either the “Lowest” or “Basement” level to create an extensive mask that included all visible Golgi signal. A cell was used for further analysis only if subtraction of the blue channel mask removed less than 65% of the area from the red channel mask. Between 75-100% of the cells typically met the criterion for further analysis. Two biological replicates (separate experiments) were performed for each quantification, as indicated by the black and blue dots on the beeswarm plots.

Representative fluorescence images chosen for display in figures were deconvolved, average projected, and scaled to the full RGB dynamic range. Then the pixel values were multiplied by 1.5 to ensure adequate visibility of both bright and faint structures. An exception is the analyses of Imh1 localization (Fig. 1 B) and Sgm1 localization (Fig. 7 C), where the images taken in the absence or presence of rapamycin were scaled identically to illustrate that the cytosolic signals in the absence of rapamycin were weak and diffuse.

### Cryo-electron tomography

Quantifoil R 2/1 grids on 300 copper mesh were prepared by glow discharging them. Log-phase yeast cultures in NSD, either mock-treated or treated for 5 min with 10 µg/ml rapamycin, were applied to the grids, which were then cryopreserved using a Vitrobot Mark IV (Thermo Fisher). The sample chamber was kept at 16 °C and 100% humidity with a 60 s wait time, and grids were blotted for 11 s before being plunged into liquid ethane. Grids were stored under liquid nitrogen.

For cryo-FIB milling, grids were clipped in AutoGrid Rings (Thermo Fisher) with a cut- out to allow for shallower milling angles. Milling was performed on an Aquilos 2 dual-beam instrument (Thermo Fisher). Samples were sputter-coated in-column with platinum for 20 s at 10 Pa and 20 mA, and then they were coated with a layer of organometallic platinum for 35 s using the gas injection system within the instrument. Targets were identified and milled using Maps software and milled using AutoTEM software (Thermo Fisher). The automated protocol included milling of micro-expansion joints at 0.5 nA, followed by rough, fine, and very fine milling at 0.5, 0.3, and 0.1 nA, respectively, followed by two polishing steps at 50 and 30 pA. The result was 10-15 lamellae per grid with a target thickness of 250-300 nm. Milled samples were stored under liquid nitrogen.

For cryo-ET and modeling, data were collected on a Titan Krios G3i at 300 kV (Thermo Fisher) with a Gatan K3 direct detection camera in CDS mode with the initial dose rate target on the detector between 7.5-8.5 electrons per pixel per second and with the Gatan BioQuantum-K3 energy slit width set to 20 eV. Tomography 5 software (Thermo Fisher) was used for collection. The tilt series was acquired with 3° steps in a bidirectional collection scheme (Hagen et al., 2017) beginning with a lamella pre-tilt of ±9° and extending to ±54°. The total dose for each tilt series was 120 e-/Å^2^. Tilts were acquired with a pixel size of 4.45 Å and with 5 µm of defocus. Tilt series data were initially reconstructed using Tomo Live software (Comet et al., 2024; Thermo Fisher), which allowed for on-the-fly motion correction, alignment, and reconstruction of tomograms using the Simultaneous Iterative Reconstruction Technique (SIRT; Gilbert, 1972). Target features were identified in the tomograms generated by Tomo Live, and those features were segmented using the 3dmod package in IMOD (Kremer et al., 1996). At least four tomograms from large budded cells were examined for each experimental condition.

To estimate the maximal distance between the plasma membrane beneath the septin ring and a tethered vesicle, we used AlphaFold 3 (https://alphafoldserver.com) to simulate the structures of FKBP-tagged Shs1, the FRB-tagged vesicle tethers, and the FRB-tagged cytosolic portions of transmembrane Golgi proteins. Assuming that the flexible protein regions were all maximally extended, the estimated lengths were 48 nm for Shs1-FKBP, 170 nm for Imh1-FRB, 68 nm for the cytosolic domain of Vps10-FRB, 39 nm for Vps52-FRB in the context of the full GARP complex (from the FRB domain to the C-terminus of Vps54), and 123 nm for Sgm1-FRB.

### Statistical analysis

For vesicle co-capture and vesicle tethering experiments, Microsoft Excel was used to compile the quantification data. Quantification of mean percentages was performed by first excluding values from cells that exhibited high levels of cisternal interference at the bud neck, where a high level was defined as removal of at least 65% of the red septin mask by subtraction of the blue Golgi mask. The data were then transferred to Prism software (Dotmatics) for plotting. Mean co-capture or tethering values were obtained for the combined individual data points from the two biological replicates. In addition, multiple linear regression was performed with a model that included two-way interactions between the biological replicate variable and the rapamycin treatment variable. The P values for the rapamycin treatment parameter (β2) are reported in the figures and legends. The P values for the replicate/treatment parameter (β3) were also examined. In some cases, this latter P value was <0.05 and the figure legend indicates that the two biological replicates were deemed to be significantly different, but those slight differences in co-capture or tethering efficiency did not affect the interpretations. Data distribution was assumed to be normal, but this assumption was not formally tested.

### Online supplemental material

Fig. S1 shows a model of the recycling pathways and tethers discussed in this paper. Fig. S2 shows supplemental data about Imh1 kinetic mapping and about the tethering of recycling proteins by ectopically localized Imh1. Fig. S3 shows cryo-ET data confirming that vesicles can be trapped by an FKBP-tagged septin at the yeast bud neck either by placing an FRB tag on the cytosolic tail of Vps10 or by ectopically localizing FRB-tagged GARP or FRB-tagged Sgm1. Fig. S4 shows supplemental data about vesicle co-capture with Vps10-FRB or Kex2- FRB. Fig. S5 shows supplemental data about vesicle tethering by ectopically localized GARP. Video 1 shows a representative 4D confocal movie of the relative arrival and departure kinetics of Imh1 and Kex2. Video 2 shows tomographic sections and modeling of part of the bud neck region in a cell with ectopically localized Imh1-FRB bound to an FKBP-tagged septin. Video 3 shows tomographic sections and modeling of part of the bud neck region in a cell with Vps10-FRB-containing vesicles captured by an FKBP-tagged septin. Video 4 shows tomographic sections and modeling of part of the bud neck region in a cell with ectopically localized Vps52-FRB bound to an FKBP-tagged septin. Video 5 shows tomographic sections and modeling of part of the bud neck region in a cell with ectopically localized Sgm1-FRB bound to an FKBP-tagged septin. Table S1 lists cryo-ET vesicle data from the tomograms of cells expressing Imh1-FRB or Vps52-FRB or Sgm1-FRB or Vps10-FRB.

## Data Availability

The data are available from the corresponding author upon reasonable request.

## Acknowledgments

This work in Benjamin Glick’s laboratory was supported by NIH grant R35 GM144050. Adam Krahn was supported by NIH training grant T32 GM007183. Thanks for assistance with fluorescence microscopy to Vytas Bindokas, Christine Labno, and Lorraine Horwitz at the Integrated Microscopy Core Facility (RRID:SCR_019197), which is supported by the NIH-funded Cancer Center Support Grant P30 CA014599. Thanks for assistance with electron microscopy to Tera Lavoie at the Electron Microscopy Core Facility (RRID:SCR_019198). We thank Luke Lavis for providing the JFX_650_ dye. Areti Pantazopoulou provided valuable comments on the manuscript. The authors declare no conflicts of interest.

## Author Contributions

Adam Krahn: conceptualization, data curation, formal analysis, investigation, methodology, validation, visualization, and writing—original draft, review, & editing. Natalie Johnson: conceptualization, data curation, formal analysis, investigation, methodology, validation, visualization, and writing—review & editing. Jotham Austin II: conceptualization, data curation, formal analysis, investigation, methodology, validation, visualization, and writing— review & editing. Benjamin Glick: conceptualization, formal analysis, funding acquisition, methodology, project administration, resources, software, supervision, validation, visualization, and writing—original draft, review, & editing.

## Abbreviations

cryo-ET: cryo-electron tomography
FKBP: FK506-binding protein
FRB: FK506-rapamycin binding domain
PBS: phosphate-buffered saline
SIRT: Simultaneous Iterative Reconstruction Technique
TGN: *trans*-Golgi network

---

**Figure S1.**
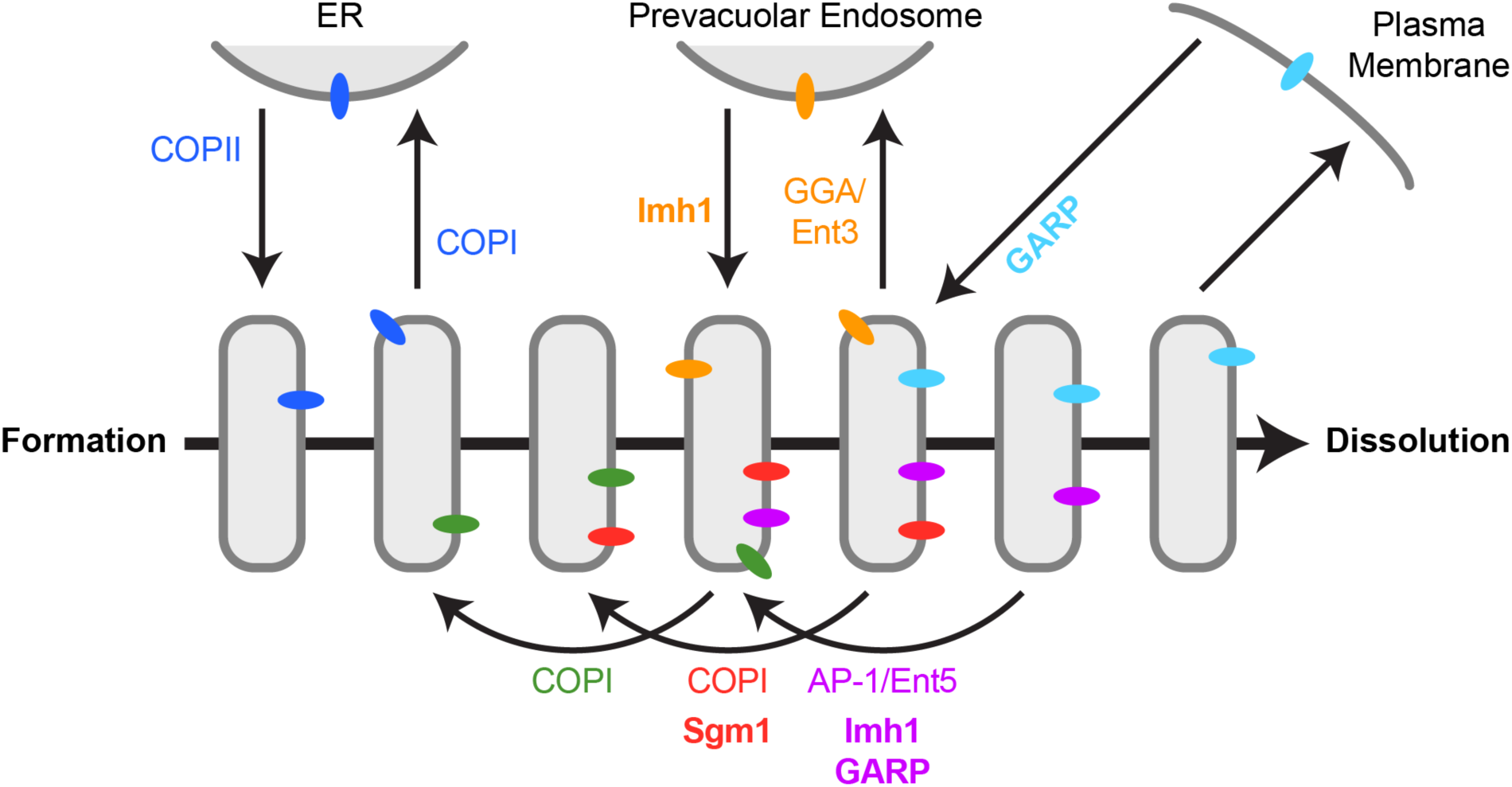
Multiple recycling pathways deliver membrane to the yeast Golgi with the aid of vesicle tethers. This diagram summarizes our interpretations about membrane traffic pathways at the yeast Golgi and the involvement of vesicle tethers. The thick arrow represents the timeline of cisternal maturation while the thin arrows represent vesicular traffic pathways. Transmembrane proteins are shown as colored ovals, and the tethers examined here are listed in bold. Six pathways are depicted. (i) COPII vesicles fuse with new Golgi cisternae, and some of the delivered proteins (dark blue) subsequently recycle to the ER in COPI vesicles. (ii) Some resident Golgi proteins such as Vrg4 (green) recycle within the Golgi in a COPI-dependent early pathway. (iii) Some resident Golgi proteins such as Tmn1 (red) recycle within the Golgi in a COPI-dependent intermediate pathway. (iv) Some resident Golgi proteins such as Kex2 (magenta) recycle within the Golgi in an AP-1/Ent5-dependent late pathway. (v) Some proteins such as Vps10 (orange) travel from Golgi cisternae to PVE compartments in a GGA/Ent3-dependent pathway and then recycle to the Golgi. (vi) Some proteins (light blue) travel to the plasma membrane in secretory vesicles and then recycle to the Golgi in endocytic vesicles. Although the endocytic pathway was not characterized in this study, GARP is presumed to tether endocytic vesicles based on evidence that GARP participates in the trafficking of endocytosed proteins (Bonifacino and Hierro, 2011) and that endocytic vesicles in yeast fuse directly with maturing Golgi cisternae (Day et al., 2018).

**Figure S2.**
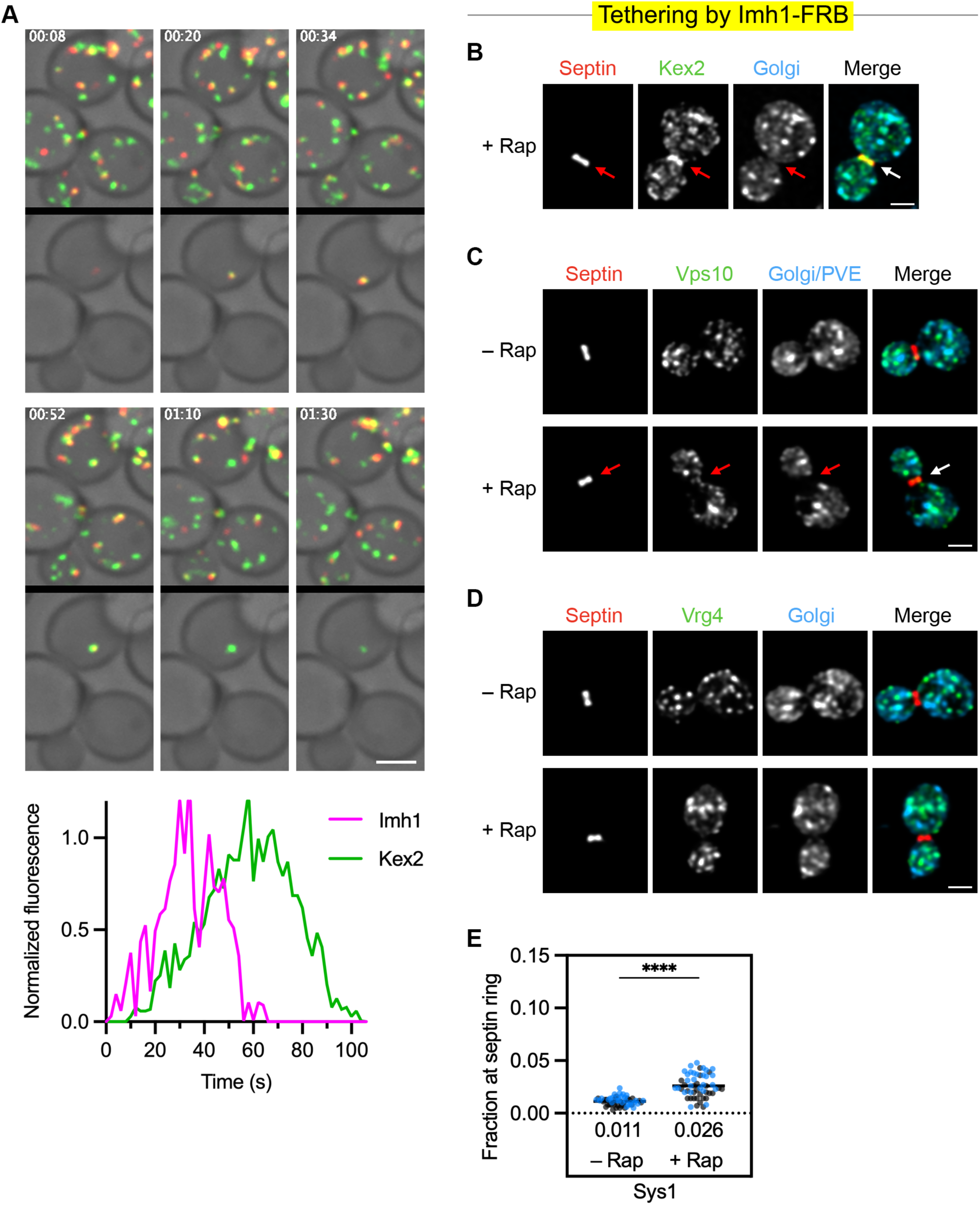
Imh1 arrives at cisternae shortly before Kex2, and ectopically localized Imh1 tethers Kex2, Vps10, and Sys1. (A) Frames from a representative 4D confocal movie of HaloTag-Imh1 (red) and Kex2-GFP (green), and kinetic traces from an individual cisterna in the movie. Average projected z-stacks are depicted for the indicated time points from Video 1. The upper row shows the complete projections, and the lower row shows edited projections that include only the cisterna that was tracked. Scale bar, 2 µm. Plotted at the bottom are normalized fluorescence intensities for the cisterna tracked in the movie. **(B)** Example of exceptionally strong tethering of Kex2 by ectopically localized Imh1 after a 10-min incubation with rapamycin (“+ Rap”). The analysis was performed as in Fig. 2 C, and this cell corresponds to the highest data point for the quantification from Fig. 2 C for the replicate shown in black. **(C)** Tethering of Vps10 by ectopically localized Imh1. Representative images show that ectopic localization of Imh1 to the FKBP-tagged septin (red) resulted in rapamycin-dependent accumulation of GFP-tagged Vps10 (green) at the bud neck. HaloTag-labeled Ric1, Sec7, and Vps8 (blue) marked Golgi cisternae and PVE compartments. These images accompany the quantification shown in Fig. 2 D. **(D)** Undetectable tethering of Vrg4 by ectopically localized Imh1. Representative images show that ectopic localization of Imh1 to the FKBP-tagged septin (red) resulted in no rapamycin-dependent accumulation of GFP-tagged Vrg4 (green) at the bud neck. HaloTag-labeled Ric1 and Sec7 (blue) marked Golgi cisternae. These images accompany the quantification shown in Fig. 2 E. **(E)** Tethering of Sys1 by ectopically localized Imh1. Fluorescence at the bud neck for GFP-tagged Sys1 was quantified with or without a 10-min rapamycin treatment as in Fig. 1 B. The total numbers of cells examined were 55 (“– Rap”) and 54 (“+ Rap”). ****, significant at P value <0.0001. Multiple regression analysis yielded a P value of 0.0275 that the two replicates were different.

**Figure S3.**
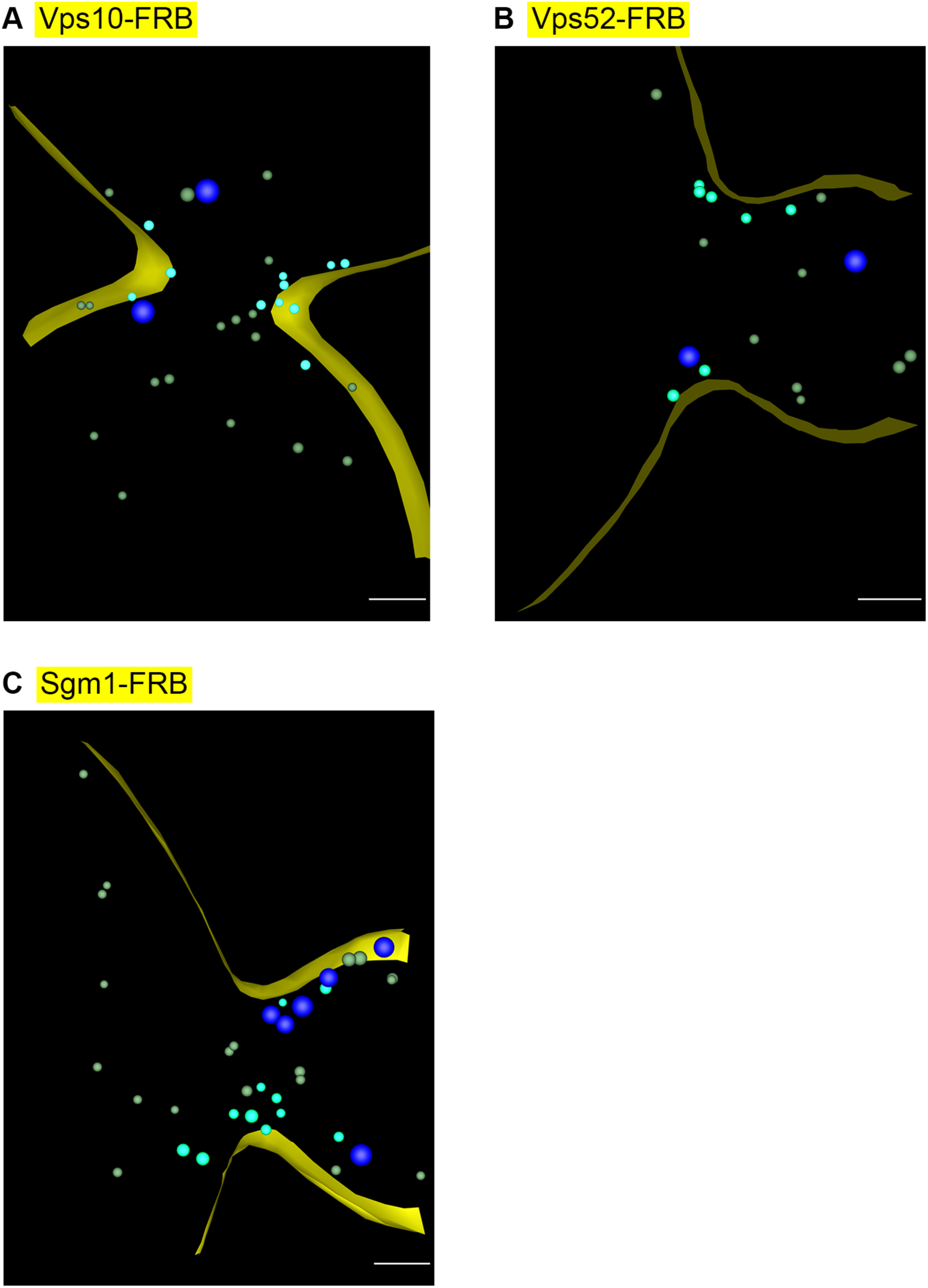
Cryo-electron tomography confirms that various classes of vesicles can be made to accumulate at the yeast bud neck using the FKBP-rapamycin-FRB system. Segmented intracellular structures are rendered as in Fig. 1 C. Scale bars, 250 nm. **(A)** Capture of vesicles carrying FRB-tagged Vps10. A log-phase culture of cells expressing Vps10-FRB and Shs1-FKBP was treated with rapamycin for 5 min followed by cryopreservation and processing for cryo-ET. Shown is the model of a SIRT-reconstructed tomogram from a large budded cell. The full data set is shown in Video 3. A vesicle was counted as putatively captured if its membrane was no more than 116 nm from a point on the plasma membrane within 200 nm from the center of the bud neck. For the 4 non-rapamycin-treated cells examined, the number of vesicles meeting this criterion ranged from 0 to 3 (mean = 1.5). For the 4 rapamycin-treated cells examined, the number of putatively captured vesicles ranged from 4 to 12 (mean = 7.0). **(B)** Tethering of vesicles by ectopically localized GARP. A log-phase culture of cells expressing Vps52-FRB and Shs1- FKBP was treated with rapamycin for 10 min followed by cryopreservation and processing for cryo-ET. Shown is the model of a SIRT-reconstructed tomogram from a large budded cell. The full data set is shown in Video 4. A vesicle was counted as putatively tethered if its membrane was no more than 87 nm from a point on the plasma membrane within 200 nm from the center of the bud neck. For the 4 non-rapamycin-treated cells examined, the number of vesicles meeting this criterion ranged from 0 to 2 (mean = 0.8). For the 4 rapamycin-treated cells examined, the number of putatively tethered vesicles ranged from 2 to 7 (mean = 4.3). **(C)** Tethering of vesicles by ectopically localized Sgm1. A log-phase culture of cells expressing Sgm1-FRB and Shs1-FKBP was treated with rapamycin for 10 min followed by cryopreservation and processing for cryo-ET. Shown is the model of a SIRT-reconstructed tomogram from a large budded cell. The full data set is shown in Video 5. A vesicle was counted as putatively tethered if its membrane was no more than 171 nm from a point on the plasma membrane within 200 nm from the center of the bud neck. For the 4 non-rapamycin-treated cells examined, the number of vesicles meeting this criterion ranged from 2 to 5 (mean = 3.3). For the 9 rapamycin-treated cells examined, the number of putatively tethered vesicles ranged from 4 to 13 (mean = 7.7).

**Figure S4.**
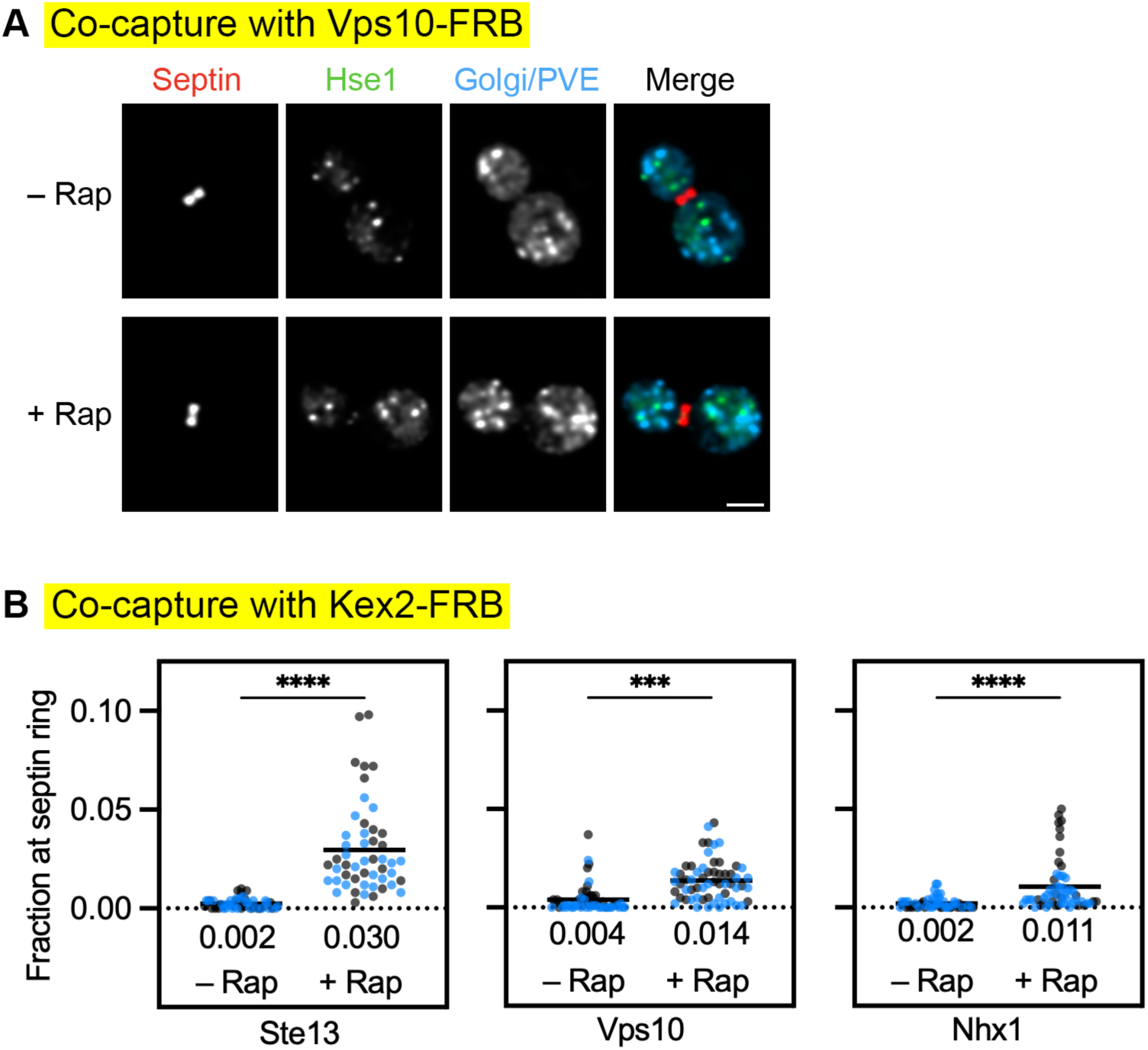
Vesicle co-capture assays can be used to examine the traffic pathways of TGN proteins. (A) Control experiment confirming minimal co-capture of the PVE marker Hse1 when Vps10-FRB is captured at the bud neck. Representative images show that capture of Vps10-FRB-containing vesicles at the FKBP-tagged septin (red) resulted in very little rapamycin-dependent accumulation of GFP-tagged Hse1 (green) at the bud neck. HaloTag-labeled Ric1, Sec7, and Vps8 (blue) marked Golgi cisternae and PVE compartments. These images accompany the quantification shown in Fig. 4 B. **(B)** Weak co-capture of GFP-tagged Vps10 or Nhx1 when Kex2-FRB is captured at the bud neck. As a control, GFP-tagged Ste13 shows relatively strong co-capture. Fluorescence at the bud neck was quantified with or without a 5-min rapamycin treatment as in Fig. 1 B. The total numbers of cells examined were 51 (Ste13 “– Rap”), 50 (Ste13 “+ Rap”), 55 (Vps10 “– Rap”), 60 (Vps10 “+ Rap”), 56 (Nhx1 “– Rap”), and 55 (Nhx1 “+ Rap”). Multiple regression analysis yielded a P value of 0.0002 that the two Nhx1 replicates were different. ****, significant at P value <0.0001; ***, significant at P value = 0.0002 (Vps10). The Ste13 co-capture data are reproduced from Krahn et al. (2026), originally published in the Journal of Cell Biology (https://doi.org/10.1083/jcb.202509174).

**Figure S5.**
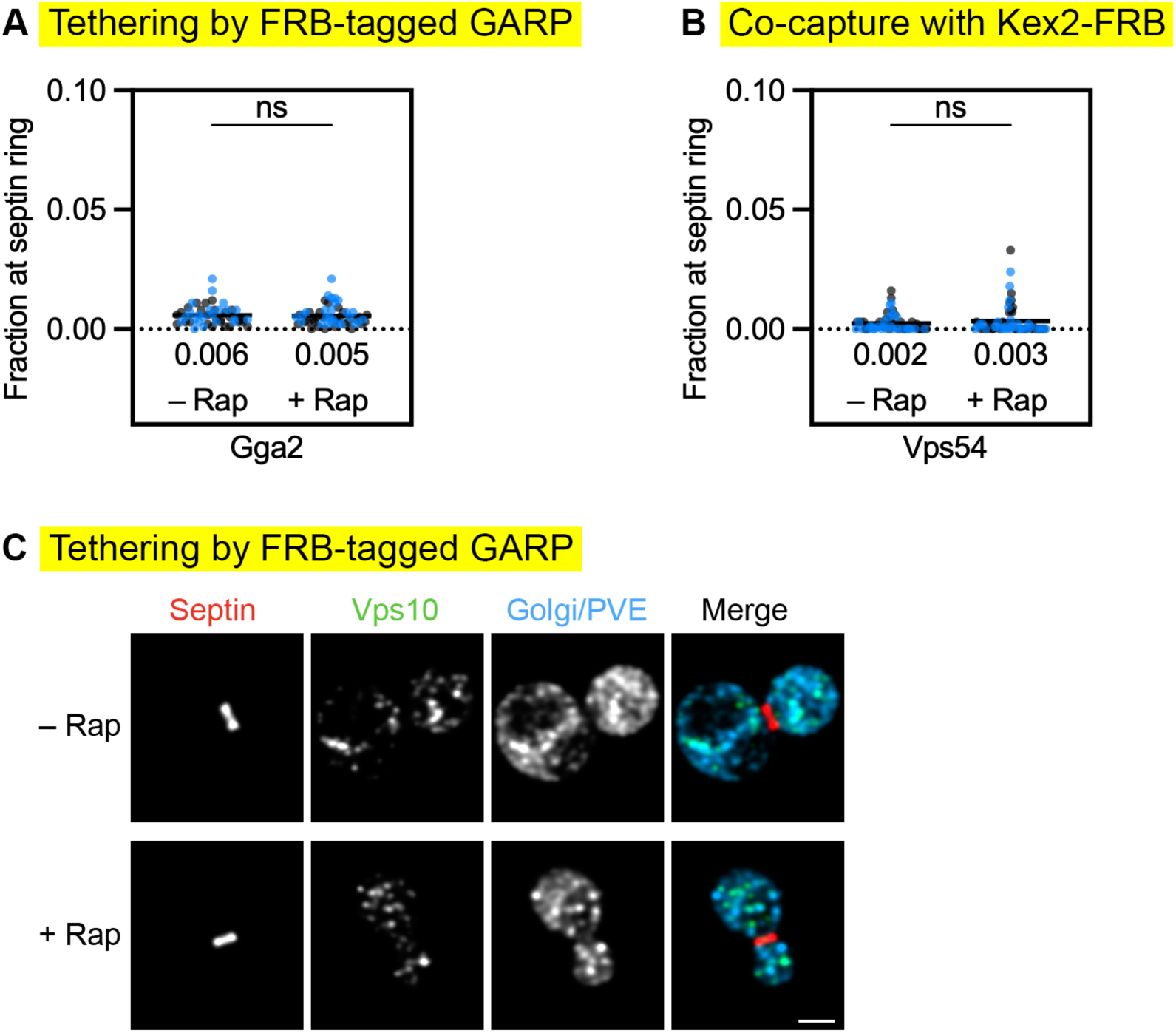
Control experiments confirm the specificity of tethering by ectopically localized GARP. (A) Control experiment showing that ectopic localization of GARP (Vps52- FRB) to the FKBP-tagged septin did not result in detectable fluorescence at the bud neck from TGN cisternae marked by Gga2. Fluorescence of GFP-tagged Gga2 at the bud neck was quantified with or without a 10-min treatment with rapamycin (“Rap”) as in Fig. 1 B. The total numbers of cells examined were 49 (“– Rap”) and 54 (“+ Rap”). ns, not significant. **(B)** Control experiment showing that capture of vesicles containing Kex2-FRB did not result in co-capture of the GARP subunit Vps54. Fluorescence of GFP-tagged Vps54 at the bud neck was quantified with or without a 5-min rapamycin treatment as in Fig. 1 B. The total numbers of cells examined were 58 (“– Rap”) and 60 (“+ Rap”). ns, not significant. **(C)** Undetectable tethering of Vps10 by ectopically localized GARP. Representative images show that ectopic localization of GARP (Vps52-FRB) to the FKBP-tagged septin (red) resulted in no rapamycin-dependent accumulation of GFP-tagged Vps10 (green) at the bud neck. HaloTag-labeled Ric1, Sec7, and Vps8 (blue) marked Golgi cisternae and PVE compartments. These images accompany the quantification shown in Fig. 6 B.

**Video 1.**
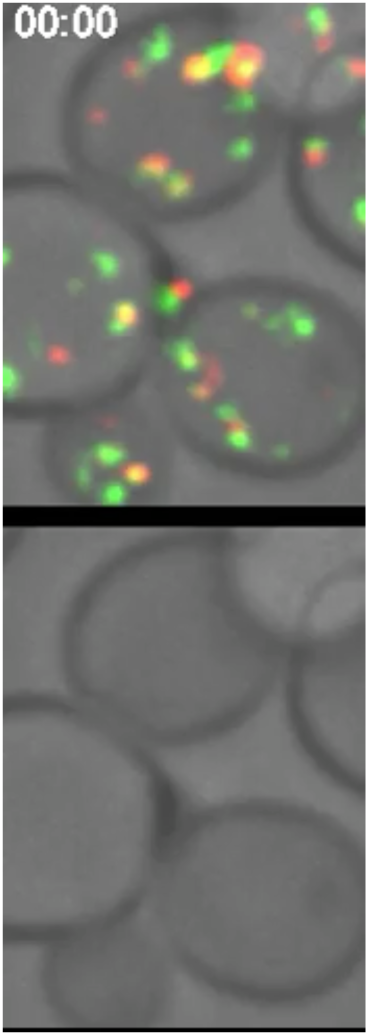
Representative 4D confocal movie of HaloTag-Imh1 and Kex2-GFP. 3D z-stacks for the individual time points were average projected. The upper row shows the complete projections, and the lower row shows edited projections that include only the cisterna that was tracked. Intervals between frames are 2 s. The overlaid numbers represent the time in seconds after the cisterna that was tracked first became detectable. See Fig. S2 A for further details.

**Video 2.**
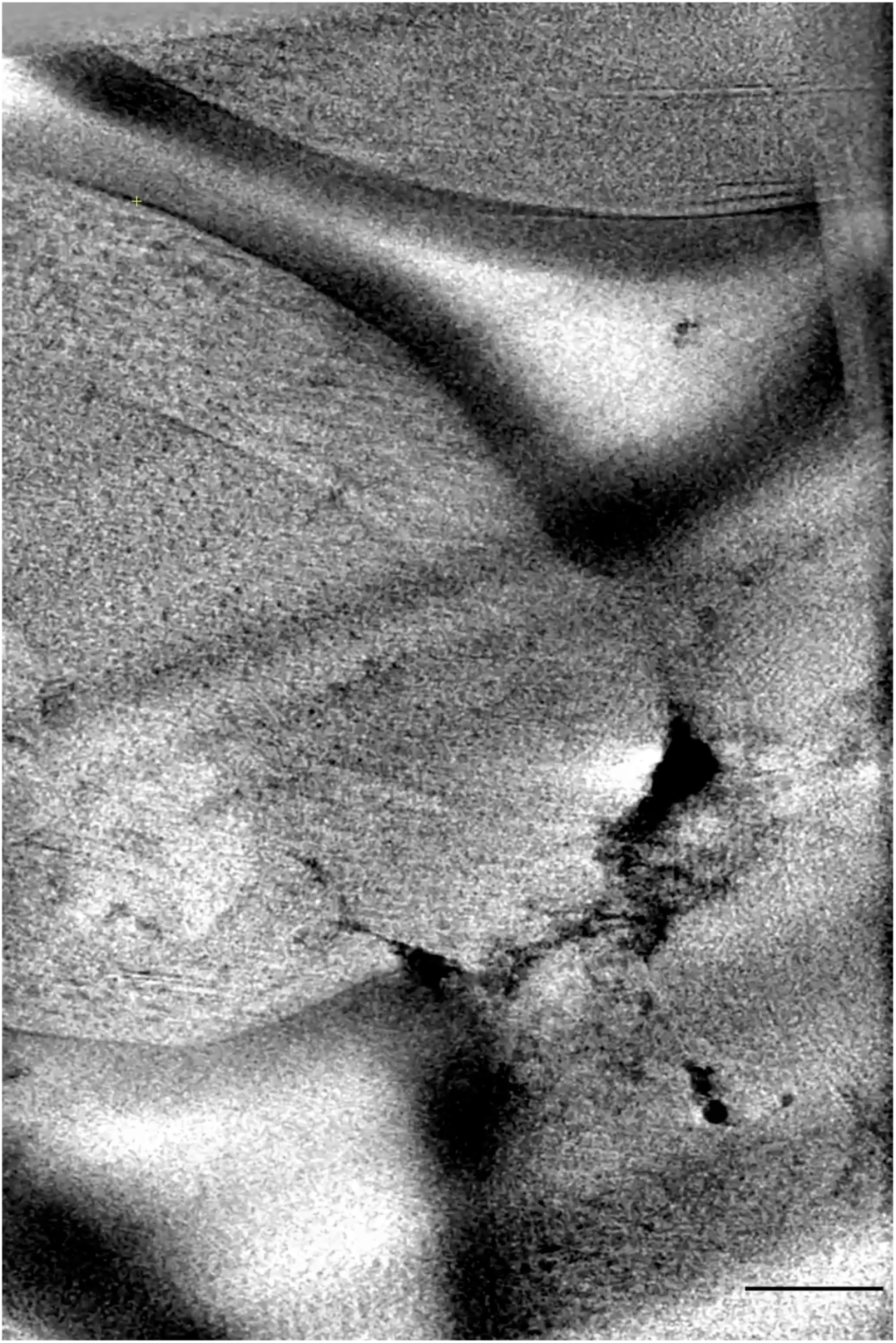
Tomographic sections and modeling of part of the bud neck region in a cell with ectopically localized Imh1-FRB bound to an FKBP-tagged septin. A log-phase culture of cells expressing Imh1-FRB and Shs1-FKBP was treated for 10 min with rapamycin prior to cryopreservation and processing for cryo-ET. The first third of the video shows every fifth section of the SIRT-reconstructed tomogram. The second third of the video shows the same tomographic sections after contours were segmented to mark the cell cortex (yellow), five secretory vesicles (blue), putatively tethered non-secretory vesicles (bright green), and other non-secretory vesicles (dull green). Also marked are a mitochondrion (cyan) and the nuclear envelope (magenta). The final third of the video shows a rotation of the tomographic model. Scale bar, 250 nm.

**Video 3.**
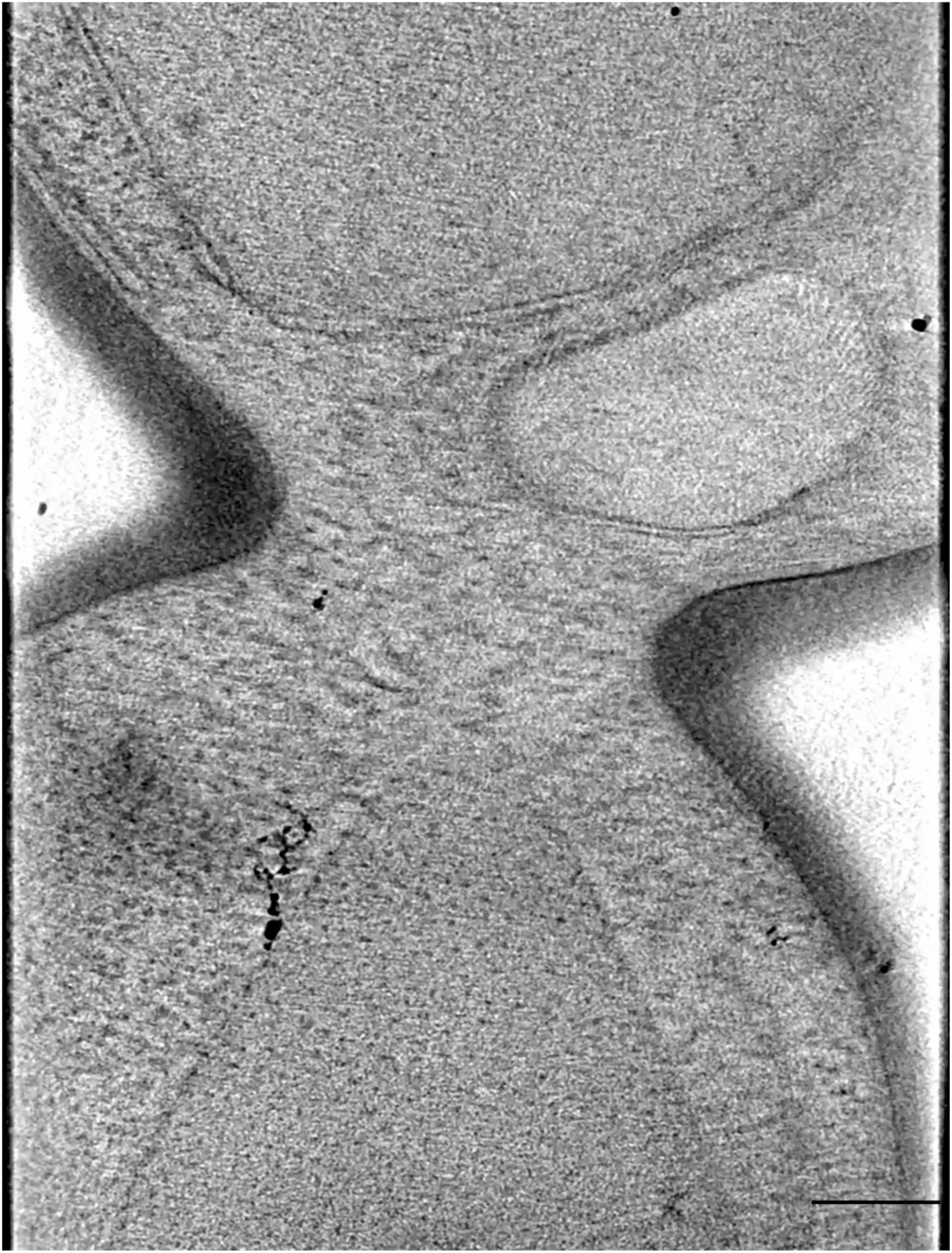
Tomographic sections and modeling of part of the bud neck region in a cell with Vps10-FRB-containing vesicles captured by an FKBP-tagged septin. A log-phase culture of cells expressing Vps10-FRB and Shs1-FKBP was treated for 5 min with rapamycin prior to cryopreservation and processing for cryo-ET. The first third of the video shows every fifth section of the SIRT-reconstructed tomogram. The second third of the video shows the same tomographic sections after contours were segmented to mark the cell cortex (yellow), two secretory vesicles (blue), putatively tethered non-secretory vesicles (bright green), and other non-secretory vesicles (dull green). Also marked are a mitochondrion (cyan) and the nuclear envelope (magenta). The final third of the video shows a rotation of the tomographic model. Scale bar, 250 nm.

**Video 4.**
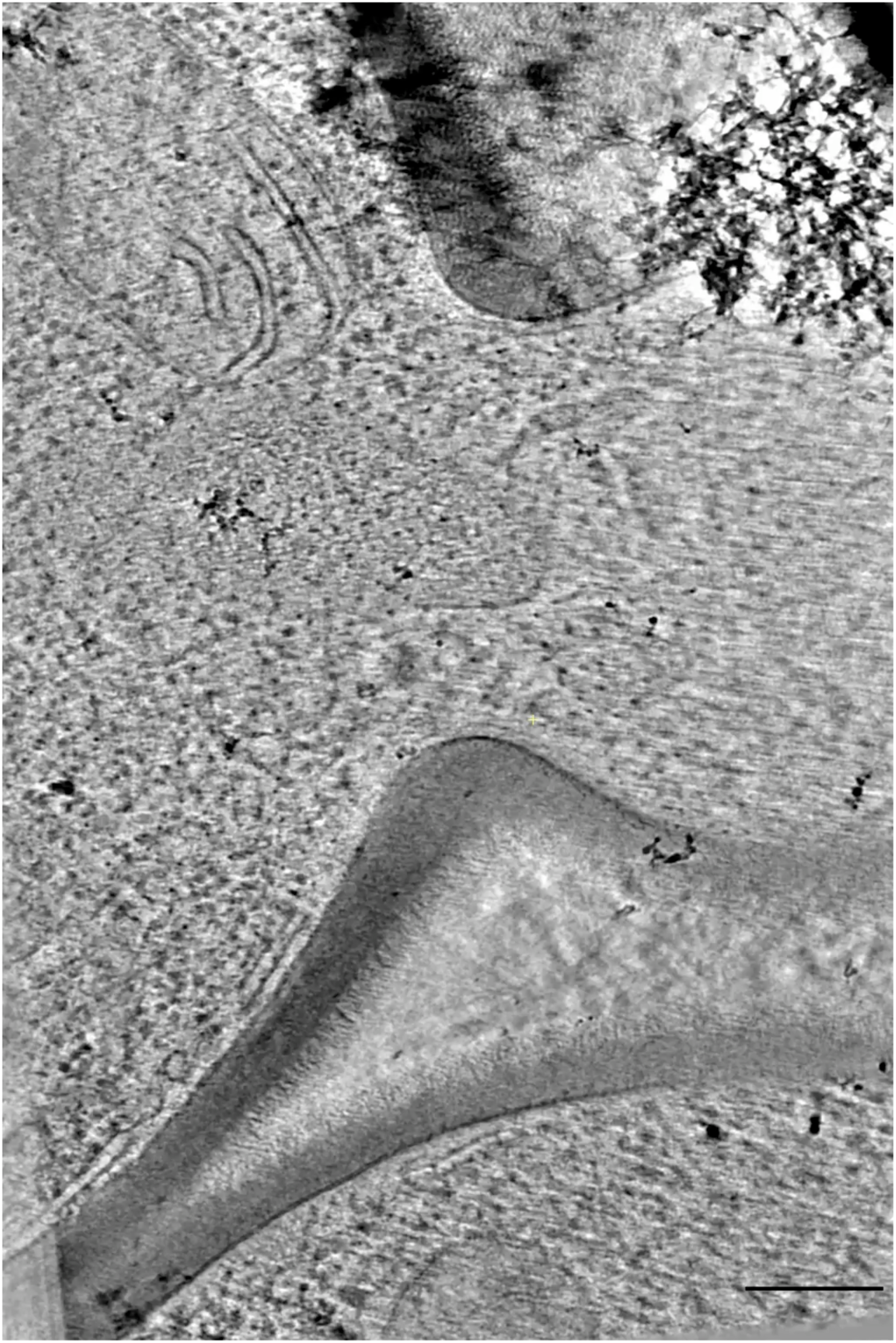
Tomographic sections and modeling of part of the bud neck region in a cell with ectopically localized Vps52-FRB bound to an FKBP-tagged septin. A log-phase culture of cells expressing Vps52-FRB and Shs1-FKBP was treated for 10 min with rapamycin prior to cryopreservation and processing for cryo-ET. The first third of the video shows every fifth section of the SIRT-reconstructed tomogram. The second third of the video shows the same tomographic sections after contours were segmented to mark the cell cortex (yellow), two secretory vesicles (blue), putatively tethered non-secretory vesicles (bright green), and other non-secretory vesicles (dull green). Also marked are two mitochondria (cyan) and the nuclear envelope (magenta). The final third of the video shows a rotation of the tomographic model. Scale bar, 250 nm.

**Video 5.**
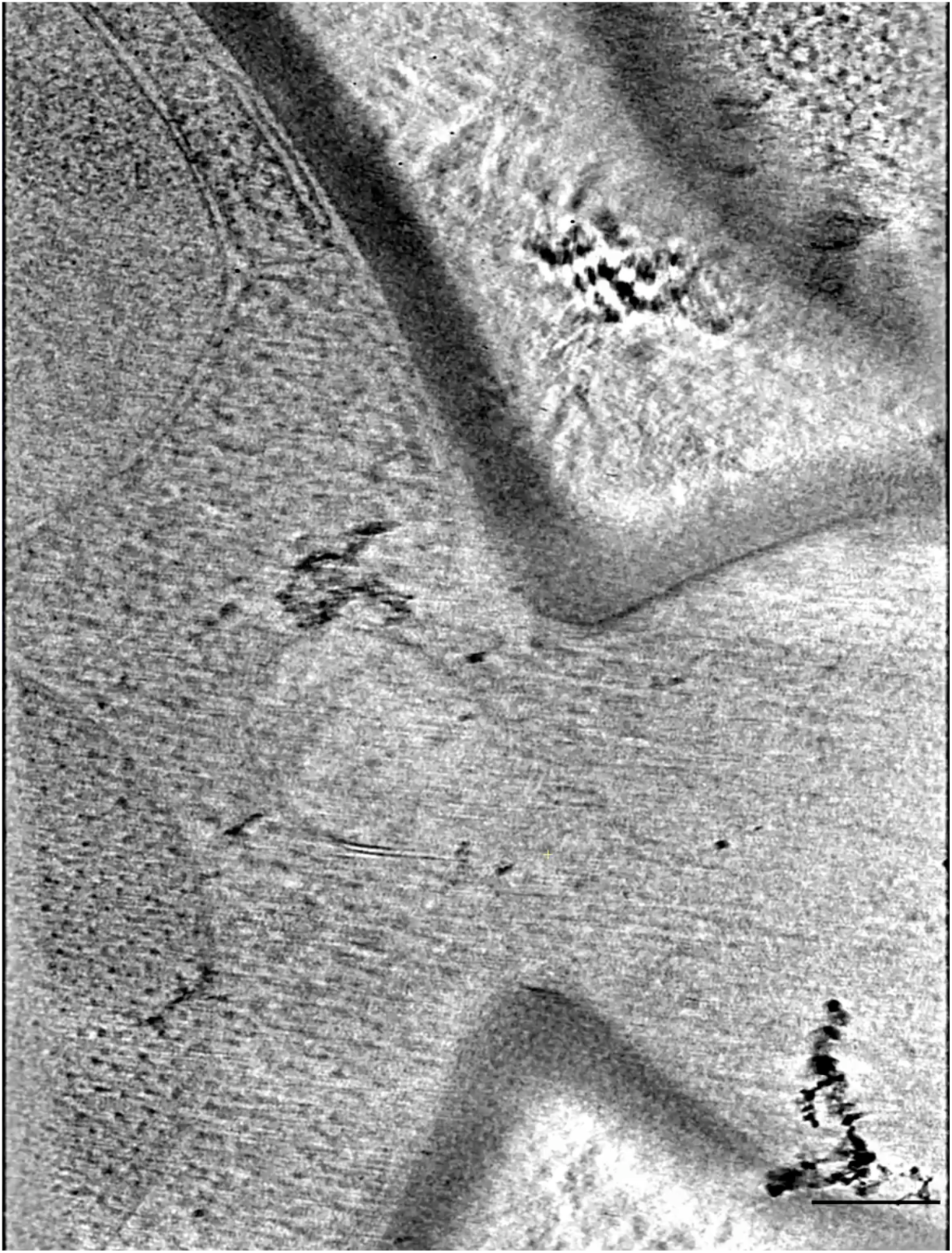
Tomographic sections and modeling of part of the bud neck region in a cell with ectopically localized Sgm1-FRB bound to an FKBP-tagged septin. A log-phase culture of cells expressing Sgm1-FRB and Shs1-FKBP was treated for 10 min with rapamycin prior to cryopreservation and processing for cryo-ET. as treated for 10 min with rapamycin prior to cryopreservation and processing for cryo-ET. The first third of the video shows every fifth section of the SIRT-reconstructed tomogram. The second third of the video shows the same tomographic sections after contours were segmented to mark the cell cortex (yellow), six secretory vesicles (blue), putatively tethered non-secretory vesicles (bright green), and other non-secretory vesicles (dull green). Also marked are a mitochondrion (cyan) and the nuclear envelope (magenta). The final third of the video shows a rotation of the tomographic model. Scale bar, 250 nm.

